# Numerosity Is Directly Sensed and Dynamically Transformed in the Human Brain: Evidence from MEG-MRI Fusion

**DOI:** 10.1101/2025.11.15.687894

**Authors:** Alireza Karami, Elisa Castaldi, Evelyn Eger, Martin Hebart, Manuela Piazza

**Author notes:** Corresponding author email address: Alireza Karami, Manuela Piazza.

## Abstract

Humans can estimate the number of objects in a scene within a fraction of a second, suggesting that numerosity is encoded rapidly and directly by the visual system. Yet how this encoding unfolds over time and interacts with other visual features remains unclear. Here, we combined magnetoencephalography (MEG) with time-resolved representational similarity analysis (RSA) and MEG–fMRI fusion to track how numerosity is represented in the brain over time. We also used multidimensional scaling (MDS) to visualize the evolving patterns of neural activity. Two main findings emerged. First, numerosity exhibited the hallmarks of a primary perceptual attribute: its neural signature appeared rapidly after stimulus onset, preceding the encoding of non-numeric features that could otherwise define number. Second, Visualization of the neural patterns using MDS suggested a temporal transformation in representational geometry, reflecting the engagement of two distinct coding schemes- an early, linear number line, consistent with a “summation code”, dominating activity in occipital regions, and a later, curved number line, consistent with “numerosity-tuned code”, emerging more strongly in associative areas along the dorsal stream. Together, these findings demonstrate that numerosity processing is encoded directly from the visual image and unfolds through a rapid hierarchical transformation, from a broad quantity signal to a finely tuned, number-specific code, linking perceptual encoding to higher-level numerical abstraction.

## Introduction

The ability to perceive numbers is widespread across the animal kingdom, from insects to humans, and is thought to have evolved due to its adaptive value: the rapid estimation of predators, prey, or food sources can be crucial for survival (Nieder, 2020). In humans, it has been suggested to act as a “start-up-system” for the development of symbolic numeracy (Piazza, 2010; Halberda et al., 2008; Decarli et al., 2023), making it a promising focus for mathematics education and remediation efforts.

The neural mechanisms by which the brain extracts numerosity from visual input remain poorly understood. One possibility is that numerosity is inferred from combinations of other quantitative, non-numeric features (e.g., item density × total field area, or total surface area ÷ average item area) (Gebuis & Reynvoet, 2012; Gebuis et al., 2016; Leibovich et al., 2016). Alternatively, numerosity may constitute a primary visual attribute, directly encoded by specialized perceptual mechanisms rather than computed from other cues (Burr & Ross, 2008; Anobile et al., 2016). Substantial behavioral evidence supports the latter view: humans and non-human primates often rely on numerosity rather than on density or total surface area when judging numerosity (Cicchini et al., 2016; Ferrigno et al., 2017), and numerosity systematically biases perception of other visual dimensions in Stroop-like tasks (Castaldi et al., 2018).

Numerosity perception is rapid and automatic, modulating physiological responses such as the pupil light reflex (Castaldi et al., 2021; Caponi et al., 2024) and driving very fast saccades, around ∼190 ms for sparse arrays versus ∼230 ms for dense, texture-like arrays (Castaldi et al., 2020).

At the neural level, fMRI studies report number-sensitive responses across occipital cortex and along both dorsal and ventral streams, with numerosity uniquely contributing to BOLD variance beyond non-numerical dimensions (Harvey et al., 2013; Castaldi et al., 2019; Karami et al., 2025). While fMRI provides precise spatial localization, its low temporal resolution leaves open whether early occipital numerosity signals reflect bottom-up encoding, feedback from higher-order regions, or independent parallel representations. The visual system comprises both feedforward and substantial feedback flows (Lamme & Roelfsema, 2000; Dehaene et al., 2006); resolving whether occipital numerosity activity initiates, receives, or independently represents numerical information therefore requires high temporal precision.

EEG and MEG are better suited to probe the timing of numerosity encoding (Gebuis & Reynvoet, 2013; Park et al., 2015; Fornaciai et al., 2017). EEG studies often report very early sensitivity to numerosity (≈75–100 ms) over posterior sensors (Park et al., 2015), supporting the idea of numerosity as a primary feature, although null results have also been reported (Gebuis & Reynvoet, 2013). However, the limited spatial specificity of EEG/MEG complicates localization: posterior scalp topographies suggest occipital involvement, but source inferences remain tentative without formal source modeling.

Frequency-tagging MEG, instead can localize feature-related responses (e.g., Van Rinsveld et al., 2021), yet it measures steady-state responses at specific frequencies and thus does not reveal the temporal sequence of neural events in the way event-related analyses do.

Another challenge in estimating numerosity is the use of composite stimulus dimensions, such as logarithmic combinations of size and spacing (e.g., Park et al., 2015; Fornaciai et al., 2017). These parameterizations are useful for mapping stimulus space but depart from perceptually meaningful features, item size, total surface area, total field area, and density, that observers can attend to and judge directly. Because different parameterizations distribute correlations among features differently, complementary tests using stimulus sets defined by perceptually relevant features are essential to uncover distinct temporal relationships among numerosity and other non-numeric features.

Overall, while past work has identified where numerosity is represented and when number-selective responses can be detected, limitations in temporal or spatial resolution, and in some cases challenges with stimulus design, leave the precise spatiotemporal dynamics underlying numerosity representations unresolved. To address this, we focus on three linked goals, which together inform our main findings: (1) to trace how numerosity and non-numerical magnitudes evolve over time (and across frequency bands); (2) to characterize how representational geometry and tuning properties of numerosity-sensitive mechanisms unfold across the cortical hierarchy; and (3) to determine where numerosity representations arise with high spatial specificity.

To achieve these aims, we combined complementary methods. First, we used time-resolved representational similarity analysis (RSA; Kriegeskorte et al., 2008) on MEG data to quantify the unique contributions of numerosity and several perceptually relevant non-numeric features simultaneously, as well as track their evolution across time and frequency. We complemented RSA with temporal generalization analysis (King & Dehaene, 2014) to test whether neural patterns encoding a feature remain stable or rapidly transform in time. Second, to visualize changes in representational geometry, we applied multidimensional scaling (MDS) to the time-resolved dissimilarity matrices, allowing us to characterize dynamic shifts in geometry and infer whether monotonic (summation-like) versus tuned coding schemes dominate at different times. Finally, to overcome the spatial ambiguity of MEG alone, we used model-based MEG–fMRI fusion via RSA (Cichy et al., 2016; Cichy & Oliva, 2020), aligning MEG timing with anatomically specific fMRI activation patterns. This multimodal framework enables a unified spatiotemporal characterization of numerosity processing. Further, it can reveal whether numerical representations emerge directly in the visual cortex, are supplied by higher areas, or arise simultaneously across regions.

By integrating these approaches, we aim to determine whether numerosity behaves as a primary visual attribute, emerging rapidly and independently of non-numeric cues, and to map how representational geometry transforms over time across cortical circuits.

## Results

Thirty healthy adult volunteers were presented with arrays of dots varying in the number of items (6, 10, 17, 29), average item areas (0.04, 0.07, 0.12, 0.2 visual square degrees), and total field areas (9 or 13.5 visual degree diameter; see Figure 1) while undergoing scanning in an MEG system. Participants were asked to estimate the number of dots and hold that information in memory until the appearance of the next set, which, in 20% of the trials, was a match stimulus, signaled by a rare change in the color of the fixation cross. Upon seeing the match stimulus, participants had to determine whether it was more or less numerous than the previous one and indicate their response by pressing a button with either their right or left hand. Participants demonstrated high overall performance (Mean=%77.81, SD=%5.83, Range=65.21% –88.54%), indicating their attentiveness and engagement in the task.

**Figure 1.**
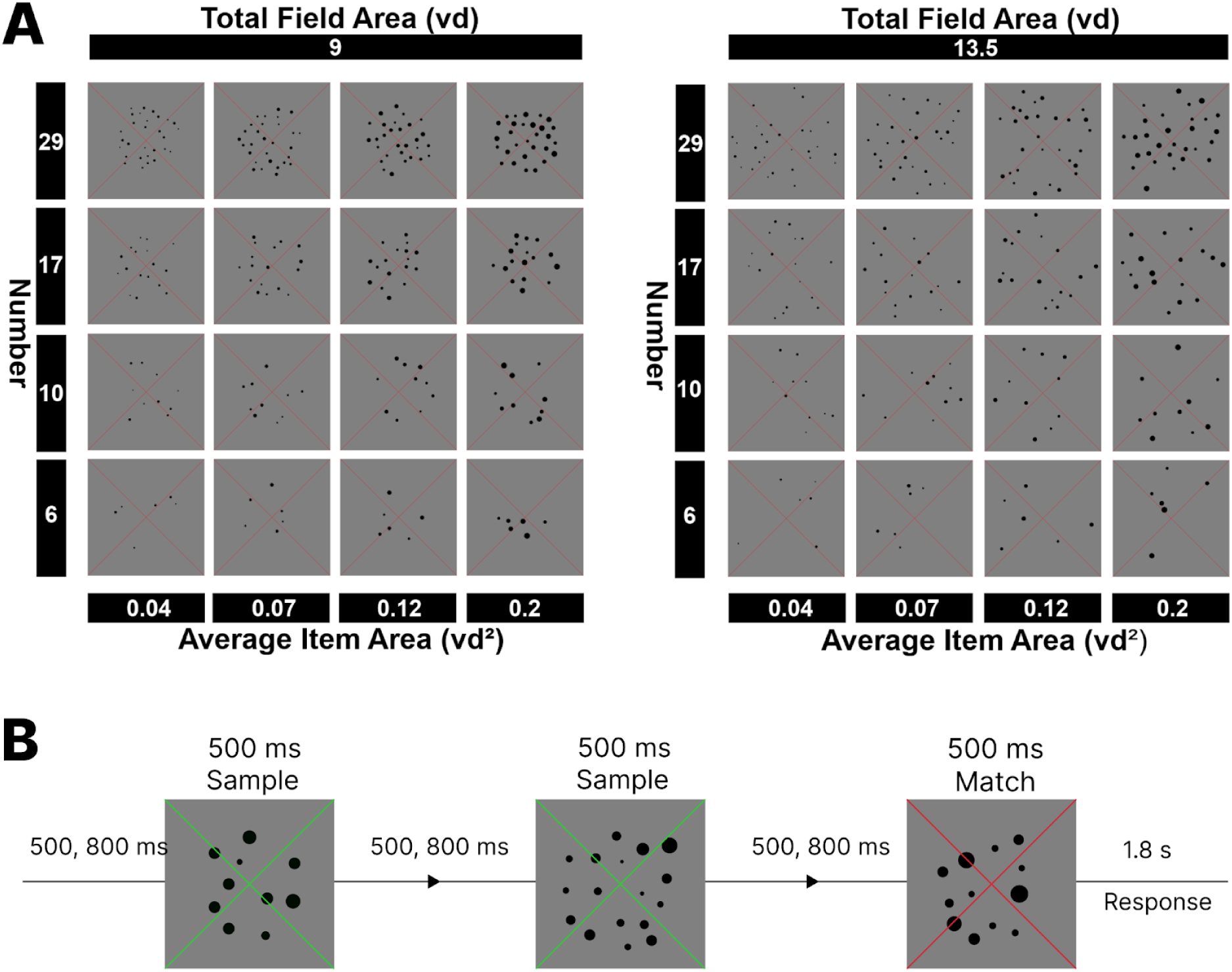
(A) A visual representation of the entire collection of stimulus conditions. The dataset encompasses orthogonal variations in multiple aspects, including the number of items (6, 10, 17, 29), the average item area (0.04, 0.07, 0.12, and 0.2 visual degree^2^), and the total field area defined by imaginary circles with diameters of 9° and 13.5°. (B) Visual representation of the chronological sequence of trials during the scanning process. Participants were instructed to focus on the number of dots and retain this information until the subsequent dot set was displayed. Periodically, the fixation cross’s color transitioned from red to green. When this color shift occurred, participants were instructed to assess whether the number of dots in the current set, known as the match stimulus, was greater or lesser than the previous one, and indicate their choice by pressing a button with their left or right hand.

### Sensor-Space Time Resolved Multivariate Representational Similarity Analysis (RSA)

In order to characterize how numeric and non-numeric feature representations change over time, we employed time-resolved multivariate RSA on the MEG signal at the sensor-space (See Figure 2). Figure 3 displays the results of the semipartial correlation between the neural and model RDMs from 100 milliseconds before the stimulus presentation to 100 milliseconds after it disappeared from the screen (−100 ms to 600 ms). Semipartial correlation ensures that the coefficients obtained reflect the variance uniquely explained by each model, while partialling out the influence of all other models. The findings revealed that the brain activity is significantly modulated by number over and above the other non-numeric features, with significant effects in three time windows: 115 – 140 ms (peak at ∼135 ms), 245 – 290 ms (peak at ∼260 ms), and 335 – 355 ms (peak at ∼340 ms) after stimulus presentation. The other non-numeric features, except average item area, showed significant effects either earlier (total field area: 65 – 600ms, peak at ∼120 ms, total surface area: 75 – 275 ms, peak at ∼165 ms) or later (density: 160 – 250, peak at ∼215 ms) than number. Importantly, while two non-numeric features (total field and total surface area) were encoded before numerosity, their specific combinations could not be used to compute number. Instead, the two feature pairs from which number could be computed (total field area and density on one side, and total surface area and average item area on the other) were not commonly encoded before numerosity. Thus, the representation of numerosity preceded that of two specific pairs of non-numeric features from which numerosity can be indirectly inferred through mathematical multiplication and division of these features, suggesting that it is directly sensed.

**Figure 2.**
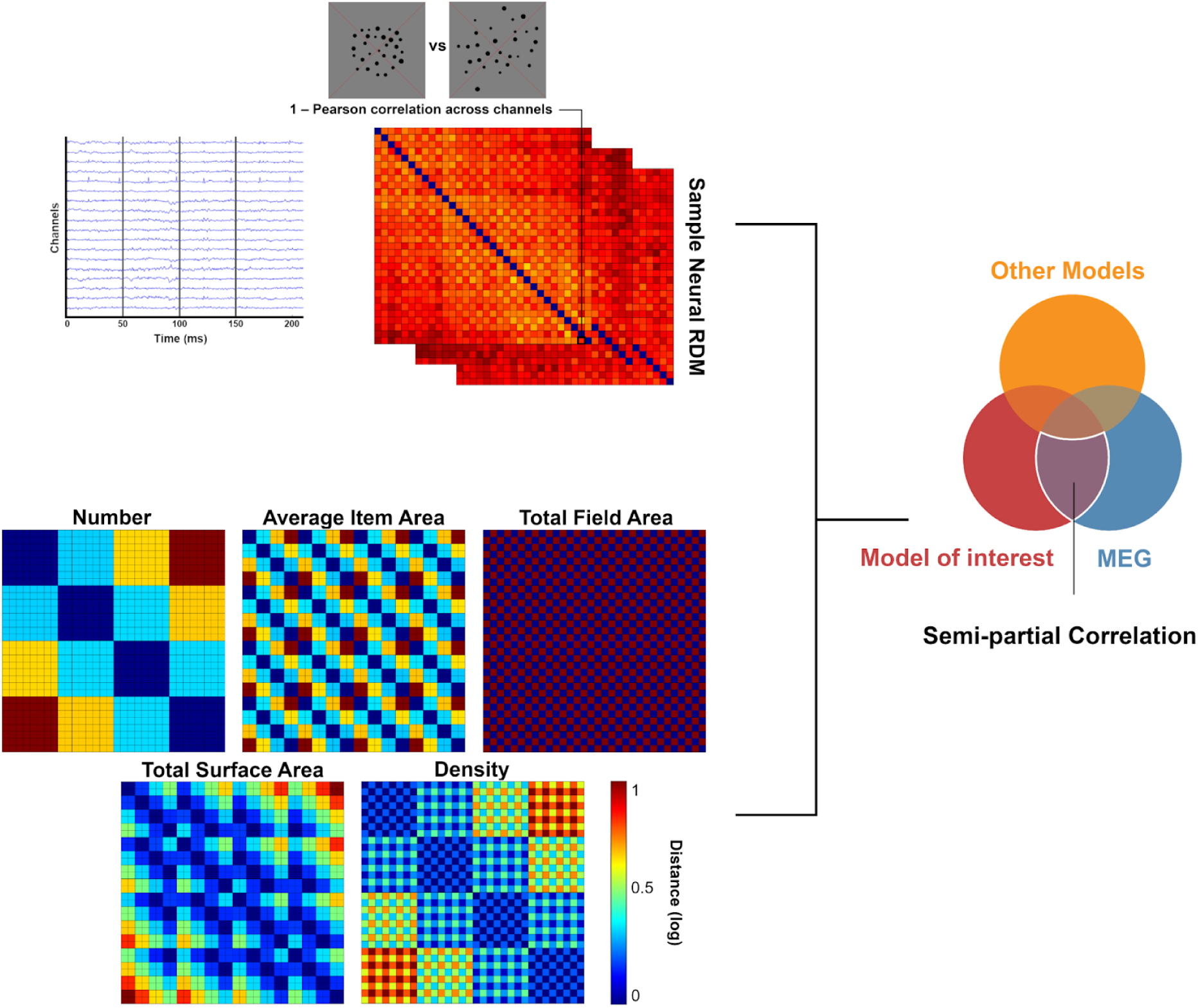
Representational dissimilarity matrices (RDMs) extracted from MEG signals were subjected to a semipartial correlation analysis. MEG RDMs were generated by computing the 1 – Pearson correlation between channel activations across time points for every pair of images. In the semipartial correlation analysis, five model RDMs were used as predictors, representing the logarithmic distance between pairs of stimuli based on number, average item area, total field area, total surface area, and density.

**Figure 3.**
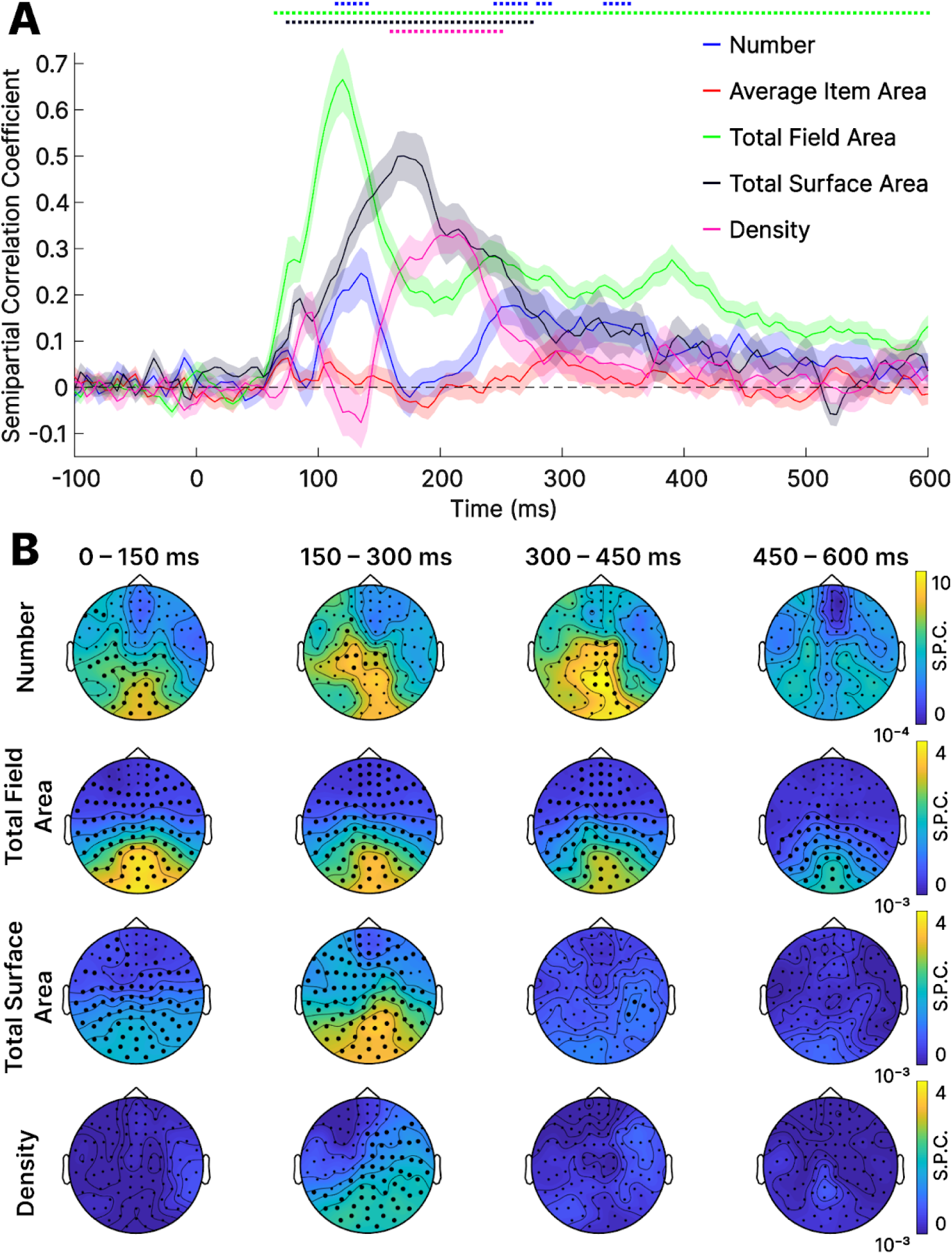
(A) Semipartial correlation (S.P.C.) coefficients derived from the representational similarity analysis for number, average item area, total field area, total surface area, and density. Standard error of the mean (SEM) across participants is depicted as a shaded area. The horizontal coloured dots indicate significant time points with effects significantly exceeding zero (thresholded at p < 0.01, TFCE corrected). (B) Each dot represents the center of a sensor-space searchlight comprising the 20 nearest MEG channels. Within each neighborhood, we performed time-resolved representational similarity analysis (RSA), computing the semi-partial correlation between the MEG RDM and each model RDM. The resulting correlation coefficients were averaged within 150-ms time bins to produce topographic maps of representational strength for each feature. The color scale indicates the semi-partial correlation coefficient (r), and bold dots mark sensors belonging to clusters showing consistent effects across participants (p < 0.01, TFCE-corrected). Group-level statistics were computed by applying cluster-based analysis across spatially adjacent sensors to identify regions of consistent effects. Average item area is not depicted here as it did not get significant. Note that the scales differ across plots in order to allow the appreciation of the effects that are smaller but significant of the non-numeric features, compared to the stronger effect of number.

### Multidimensional Scaling

Second, to visualize the evolving structure of the neural representational space we applied multidimensional scaling (MDS) to the group-average representational dissimilarity matrices (RDMs) computed across thirty participants. MDS provides a low-dimensional representation that makes the geometry of pairwise similarities interpretable. We computed MDS solutions at successive time points from stimulus onset up to 600 ms, including a 100 ms post-offset window (see Videos 5–7).

The resulting MDS reveal a clear rank-ordering of numerosities emerging around ∼70 ms after stimulus onset. Later, between approximately 220 and 315 ms, the low-dimensional configuration departs from a linear arrangement and displays a pronounced curved geometry (for snapshots see Figure S2 in supplementary material). This curvature indicates a reorganization of the representational geometry and closely mirrors the parietal manifold we reported previously in fMRI (Karami et al., 2025), as well as recent MEG findings using a different stimulus range (Barnett & Fleming, 2024). Consistent with the interpretation proposed by Karami et al. (2025) based on simulation results, the early linear MDS pattern may reflect a monotonic code (cells which responses scale with numerosity), whereas the later curved manifold likely corresponds to numerosity-tuned populations (cells selectively tuned to specific numbers).

### Sensor-Space Searchlight RSA

Results from the sensor-space searchlight semi-partial correlation RSA (Figure 3) indicate that information related to number independent from other features was present in the MEG channels positioned over the occipital and parietal cortex already within 150 ms from stimulus presentation and was further amplified in time up to 450 ms. In contrast, non-numeric features showed significant contributions mainly over occipital sensors, with maximal effects in the 100–250 ms range that progressively declined after ∼300 ms. In terms of timing, the observation that pure number information peaked in the MEG channels beyond the strictly posterior ones within 150 ms confirmed that numerical representations emerge very early, suggesting that they are present also in regions that extend beyond the primary visual cortex.

### Time-Frequency Resolved RSA

We then further expanded the analysis from the time domain to the time-frequency domain. The rationale behind this approach lies in the possibility that if numbers are represented in a distinct frequency range compared to other non-numeric features, it suggests that numeric and non-numeric features are represented through different functional networks. Our time-frequency analysis, depicted in Figure 4, demonstrated that number is represented independently from other visual features in the lower beta range (12-17 Hz), whereas density and total surface area are represented in the higher beta range (17-30 Hz). Total field area is represented in a broader frequency range that encompasses both lower and higher beta. However, the total field area varied orthogonally to number in our experiment. Thus the fact that it is represented in the same frequencies cannot be taken as an index for their functional dependency. Finally, the average item area did not show any statistically significant effects in any frequency band (thresholded at p < 0.01, TFCE corrected).

**Figure 4.**
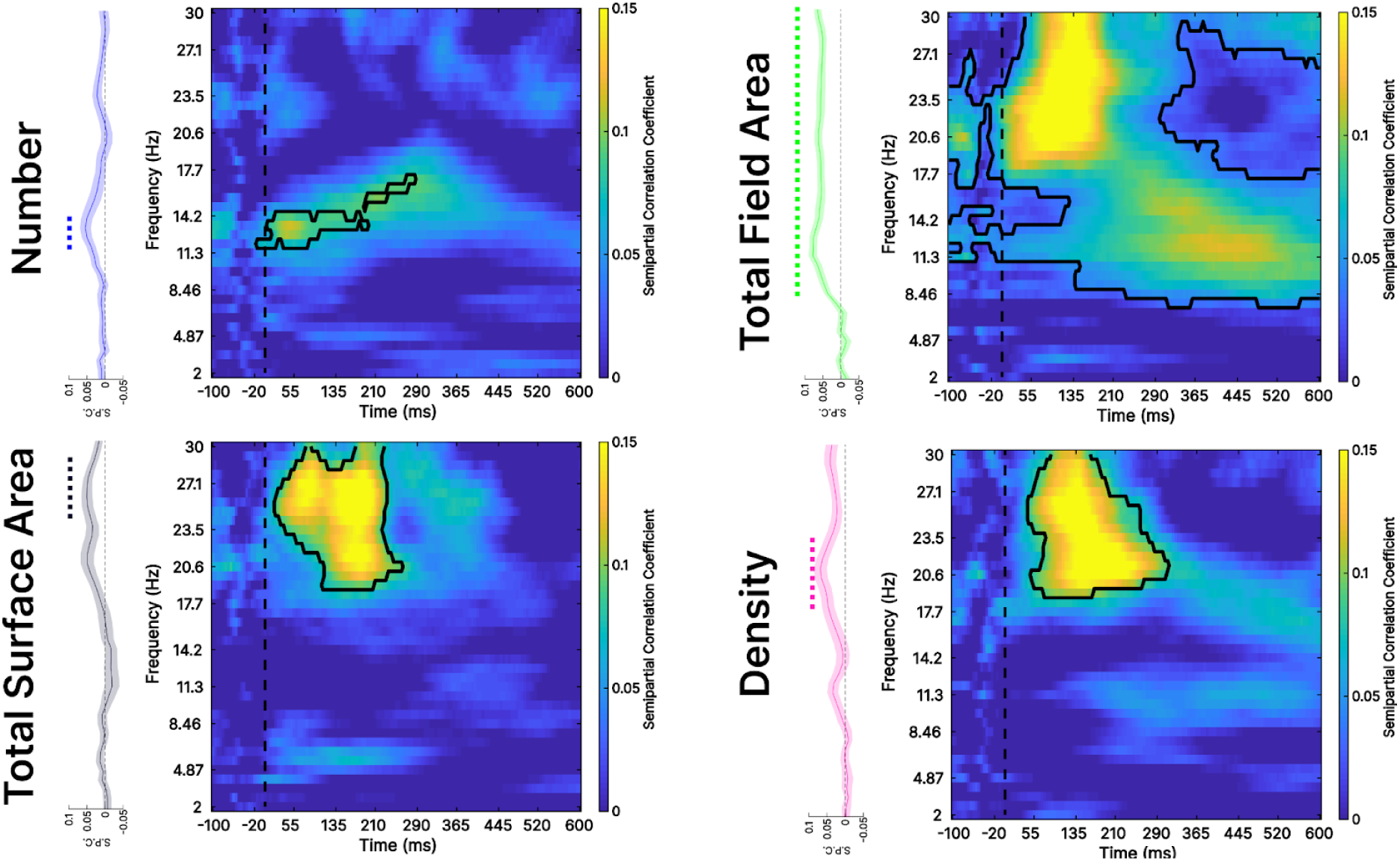
Semipartial correlation coefficient obtained from time-frequency resolved RSA for number, average item area, total surface area, and density. The black outline reflects the significant clusters (thresholded at p < 0.01, TFCE corrected) and the dashed line represents time zero. The vertical figure next to each matrix represents the average of individual frequencies across time. The colored dots above them indicate significant frequencies (thresholded at p < 0.01, TFCE corrected). Average item area is not depicted here as it did not get significant.

### Temporal Generalization Analysis

To test the temporal stability of numeric and non-numeric feature representations, we ran a temporal-generalization analysis (King & Dehaene, 2014). Classifiers trained at one timepoint generalized broadly across time for numerosity and three non-numeric features (total field area, total surface area, and density), yielding off-diagonal decoding patterns consistent with a stationary, time-stable representational format. In contrast, average item area produced a pronounced thick diagonal in the temporal-generalization matrix (Figure 5): decoding was reliable only when training and testing involved the same or neighboring time windows and failed to generalize to distant timepoints. This diagonal profile indicates a rapidly transforming, time-specific code for average item area, with the discriminative neural pattern shifting over the course of processing rather than remaining fixed.

**Figure 5.**
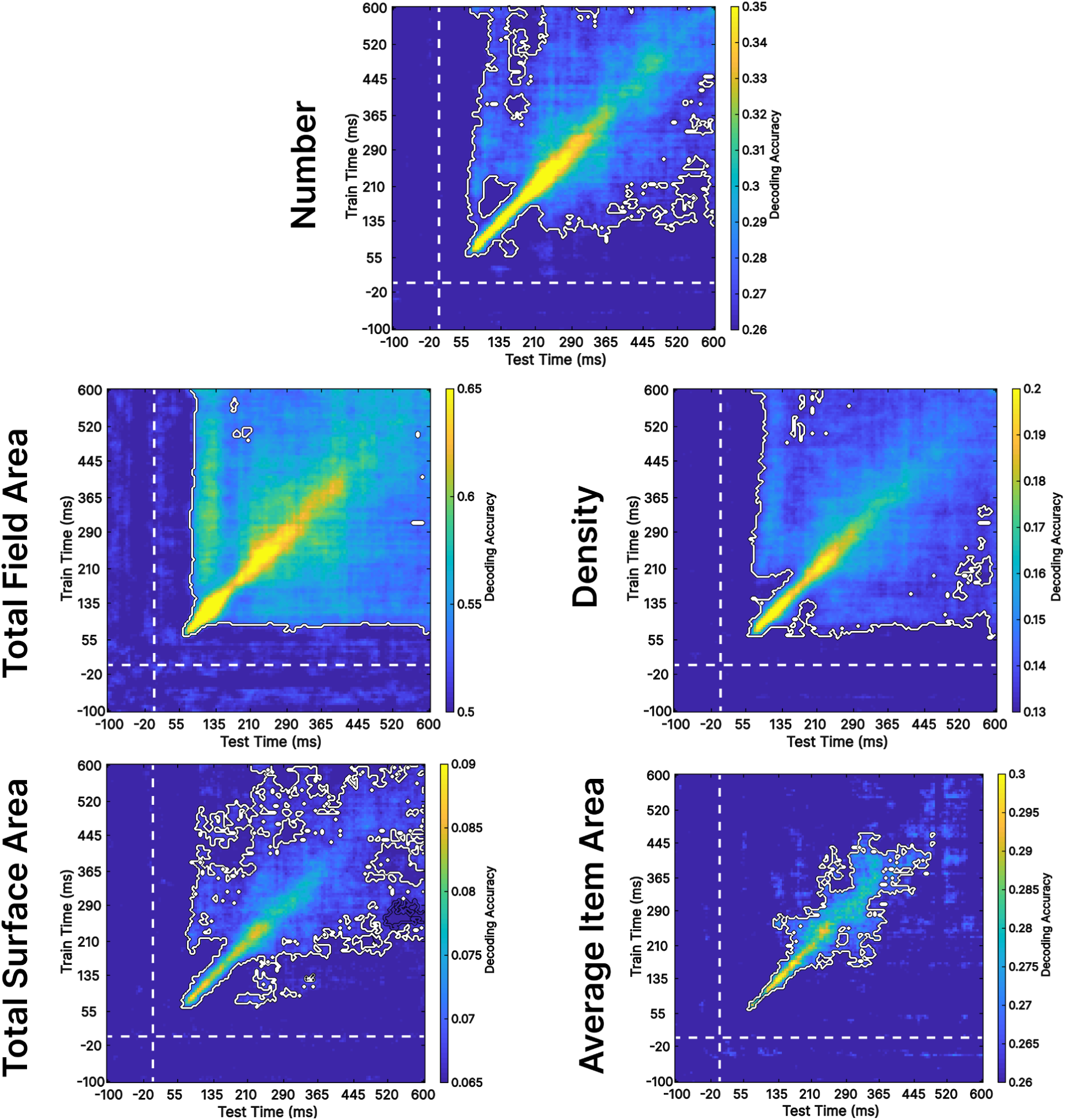
The temporal generalization matrix reflects the decoding accuracy (with a chance level of 0.25 for number, 0.5 for total field area, 0.0625 for total surface area, and 0.125 for density). The y-axis represents the training time during which the classifier was trained to classify among four numbers (6, 10, 17, 29), two total field areas (9, 13.5), sixteen total surface areas (resulting from multiplying four different numbers by four distinct average item areas), and eight different densities (resulting from dividing four different numbers by two different total field areas). Meanwhile, the x-axis represents the test time when the classifier was tested to decode the numbers, total field area, total surface area, and density. The white outline reflects the significant clusters (thresholded at p < 0.01, TFCE corrected) and the dashed line represents time zero.

The motivation for this analysis is straightforward: if numerosity (or any other feature) is encoded in a time-stable format it can be read out by downstream regions using the same population code across an extended interval. Conversely, a dynamic format implies rapid reconfiguration of neural patterns. Such short-lived, potentially sign-reversing activity may be obscured in fMRI due to temporal and spatial averaging (Rogers et al., 2021), which could explain why analogous representations (e.g., for average item area) were not detected in previous fMRI studies (Castaldi et al., 2019; Karami et al., 2025).

### MEG-fMRI Fusion with RSA

To combine the MEG data from the current experiment with the fMRI data from our previous study (Karami et al., 2025) we then applied the MEG–fMRI fusion approach. Both datasets were collected using the same task and experimental design, as detailed in the *Materials and Methods* section, with the only difference being the faster stimulus presentation rate in the MEG experiment. The fusion analysis was conducted across time for the MEG data and across predefined regions of interest (ROIs) derived from fMRI. Specifically, in line with our previous fMRI analyses, we examined three regions along the dorsal stream (V1–V3, V3ABV7, and IPS1–5) and three regions along the ventral stream (V1–V3, VO, and PHC).

Across both processing streams, neural representational correspondence between MEG and fMRI began to emerge around 60 ms after stimulus onset. Along the dorsal stream, the average peak latencies across participants were 172.50 ms (SD = 104.74) for V1–V3, 183.33 ms (SD = 52.71) for V3ABV7, and 206.50 ms (SD = 49.83) for IPS1–5. To statistically assess these latency differences, we performed Wilcoxon signed-rank tests comparing peak times between regions. The results revealed that IPS1–5 peaked significantly later than V1–V3 (*p* = 0.0014). In contrast, the differences between V1–V3 and V3ABV7 (*p* = 0.0136) and between V3ABV7 and IPS1–5 (*p* = 0.0562) did not reach significance.

### Model-based MEG-fMRI Fusion

Prior fMRI studies have demonstrated that numerical information is represented across retinotopic regions in both the dorsal stream up to the Intraparietal Sulcus (Castaldi et al., 2019; Karami et al., 2025) and the ventral stream up to Ventral Occipital (Karami et al., 2025). Results from the time-resolved RSA in the previous sections indicate that pure numerical information, independent of other non-numeric features, is represented very early in time. However, due to the limited spatial resolution of MEG and the sluggish nature of BOLD signals in fMRI, a detailed understanding of brain activity across time in individual regions of interest remains elusive. For example, the representation of numerosity observed in early visual cortex in the prior fMRI studies (Castaldi et al., 2019; Karami et al., 2025) might either reflect a feedforward processing starting there but it might also be the result of feedback signals from higher-order cortical areas. To distinguish between these hypotheses, we employed a model-based MEG-fMRI fusion approach (Khaligh-Razavi et al., 2018; Hebart et al., 2018). In its present format, the model-based fusion of MEG and fMRI involves the conjunction on significant results from three distinct representational similarity analyses: MEG-to-fMRI RSA (see Figure 6), MEG-to-model (see Figure 2), and fMRI-to-model RSA (See Figure S1 in the supplementary material; see also Karami et al., 2025) for five different model RDMs, namely number, average item area, total field area, total surface area, and density. The findings from the model-based MEG-fMRI fusion is shown in Videos 1 to 4 (with temporal resolution of 5 milliseconds) for number, total field area, total surface area, and density respectively (for a snapshot of each feature in the dorsal stream regions see Figure 7 and the ventral regions see Figure 8). The results showed that pure numerical information was equally rapidly represented across regions in the early visual area (V1-3), dorsal (V3ABV7, and IPS1-5) and ventral (VO, PHC) streams, preceding pairs of non-numeric features (total field area and density, or total surface area and average item area) which could indirectly contribute to the encoding of numerical information. The fact that we were not able to detect variations in the onset of numeric and non-numeric features across different regions, suggests the parallel emergence of these representations in these brain regions within the temporal precision afforded by our analysis approach. Together, these results highlight a direct and fast representation of numerical information along both the dorsal and ventral streams regions.

**Figure 6.**
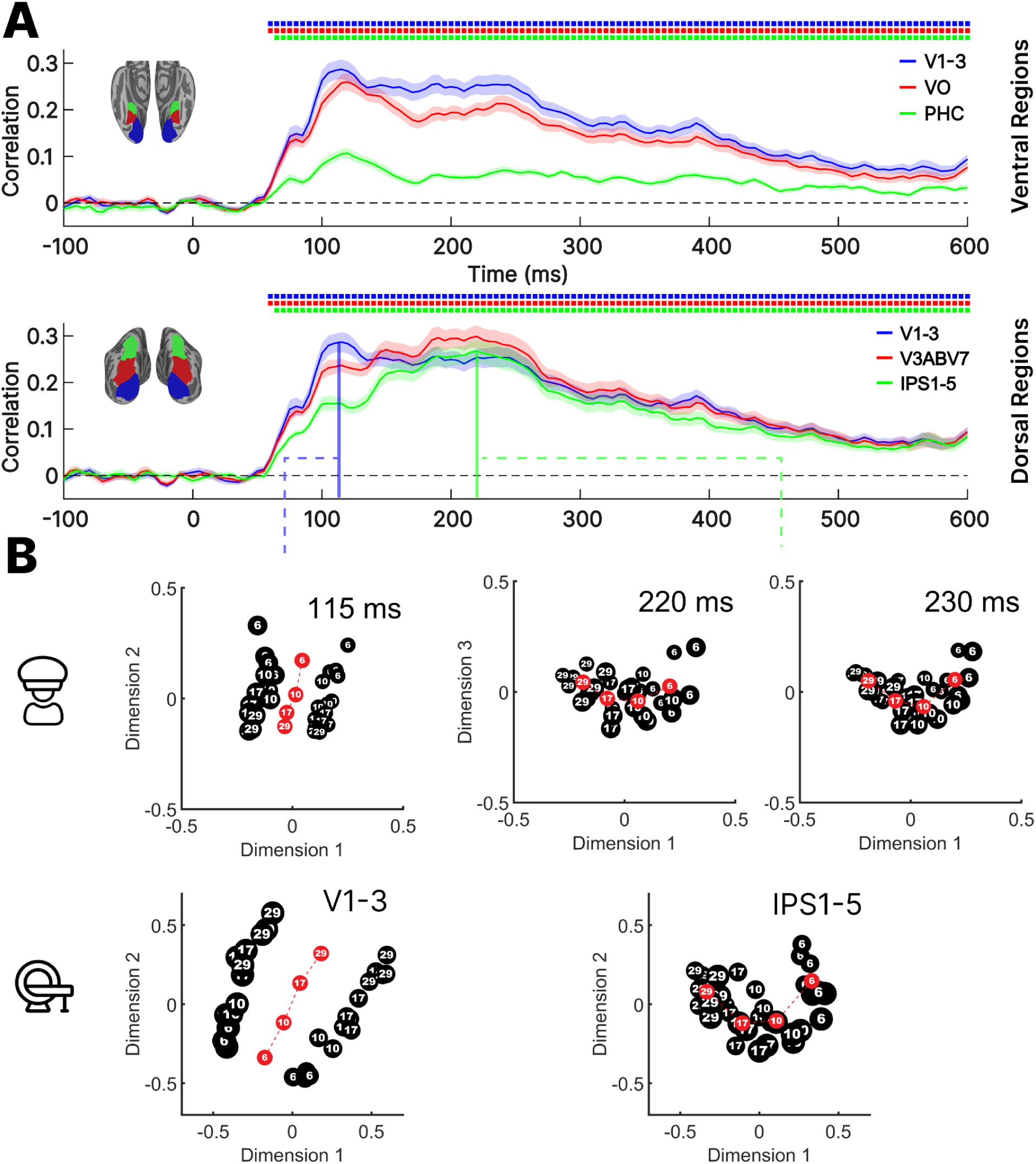
(A) Pearson correlation coefficients from the representational similarity analysis between individual participants’ MEG RDMs and group-averaged fMRI RDMs (N = 31) are shown for early visual areas (V1–V3), dorsal regions (V3ABV7, IPS1–5), and ventral regions (VO, PHC). The shaded areas represent the standard error of the mean (SEM) across participants. Colored horizontal dots mark time points with significant correlations exceeding zero (p < 0.01, TFCE-corrected). (B) Multidimensional scaling (MDS) illustrates representational similarities between stimuli in a two-dimensional space at three key time points corresponding to the peaks observed in the group-averaged time series across regions: the peak in V1–V3 at 115 ms, the peak in IPS1–5 at 220 ms, and a subsequent point at 230 ms. The second row shows corresponding MDS results from our previous fMRI study in V1–V3 and IPS1–5. The black circles represent the 32 stimuli. The circle sizes vary, indicating stimuli with small total field area (small circles) and larger total field area (large circles). The red circles indicate the average coordinates of each number.

**Figure 7.**
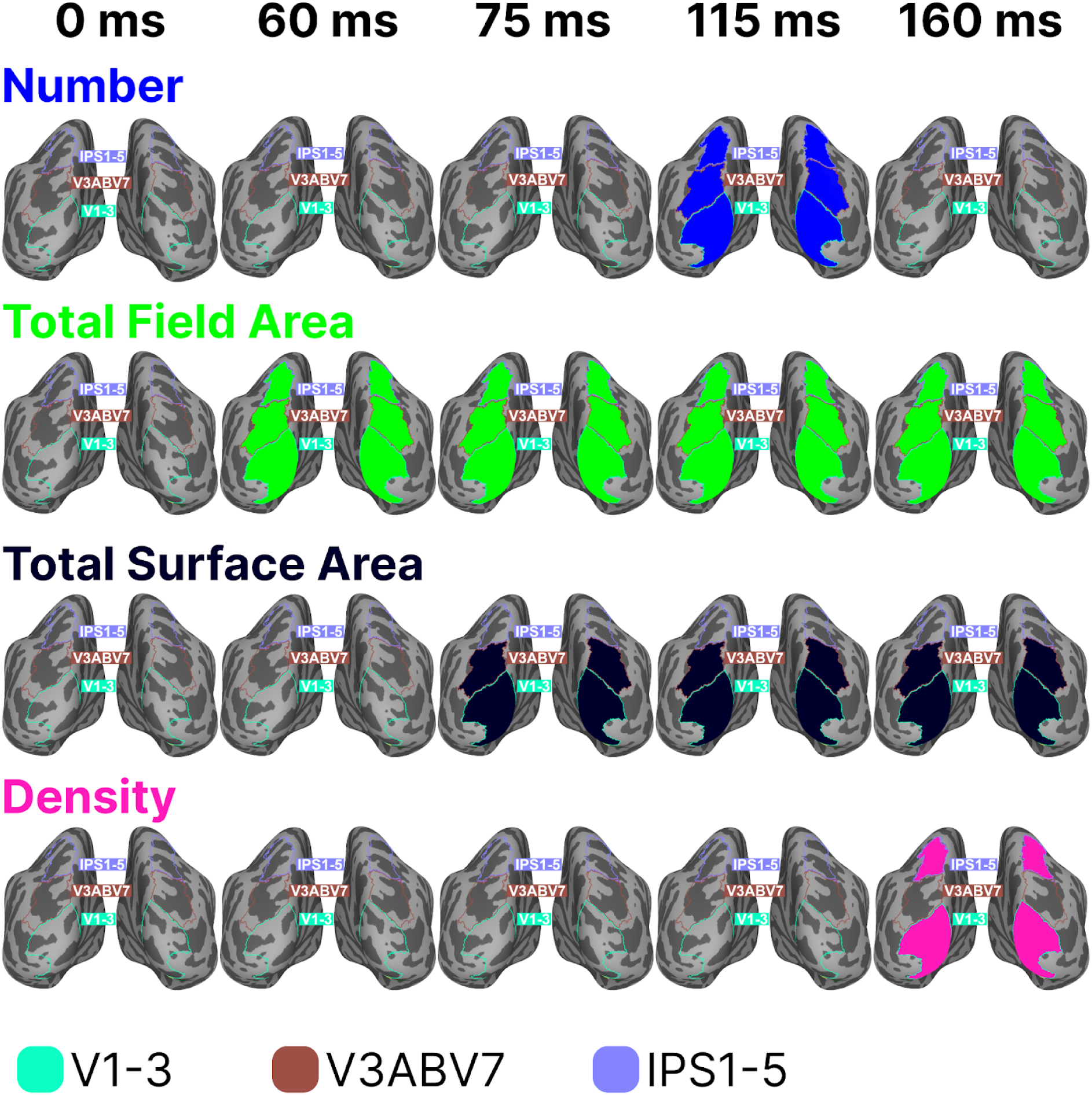
Spatiotemporal dynamics of four features (number, total field area, total surface area, and density) in the dorsal stream regions are shown at five snapshots (time 0 ms when the stimulus begins, time 60 ms when total field area becomes significant, time 75 ms when total surface area becomes significant, time 115 ms when number becomes significant, and 160 ms when density becomes significant) from Videos 1 to 4. It’s important to note that the average item area is not depicted, as it did not show significance in any region. For a higher temporal resolution (5 milliseconds) visualization of the results, please refer to supplementary videos 1 to 4)

**Figure 8.**
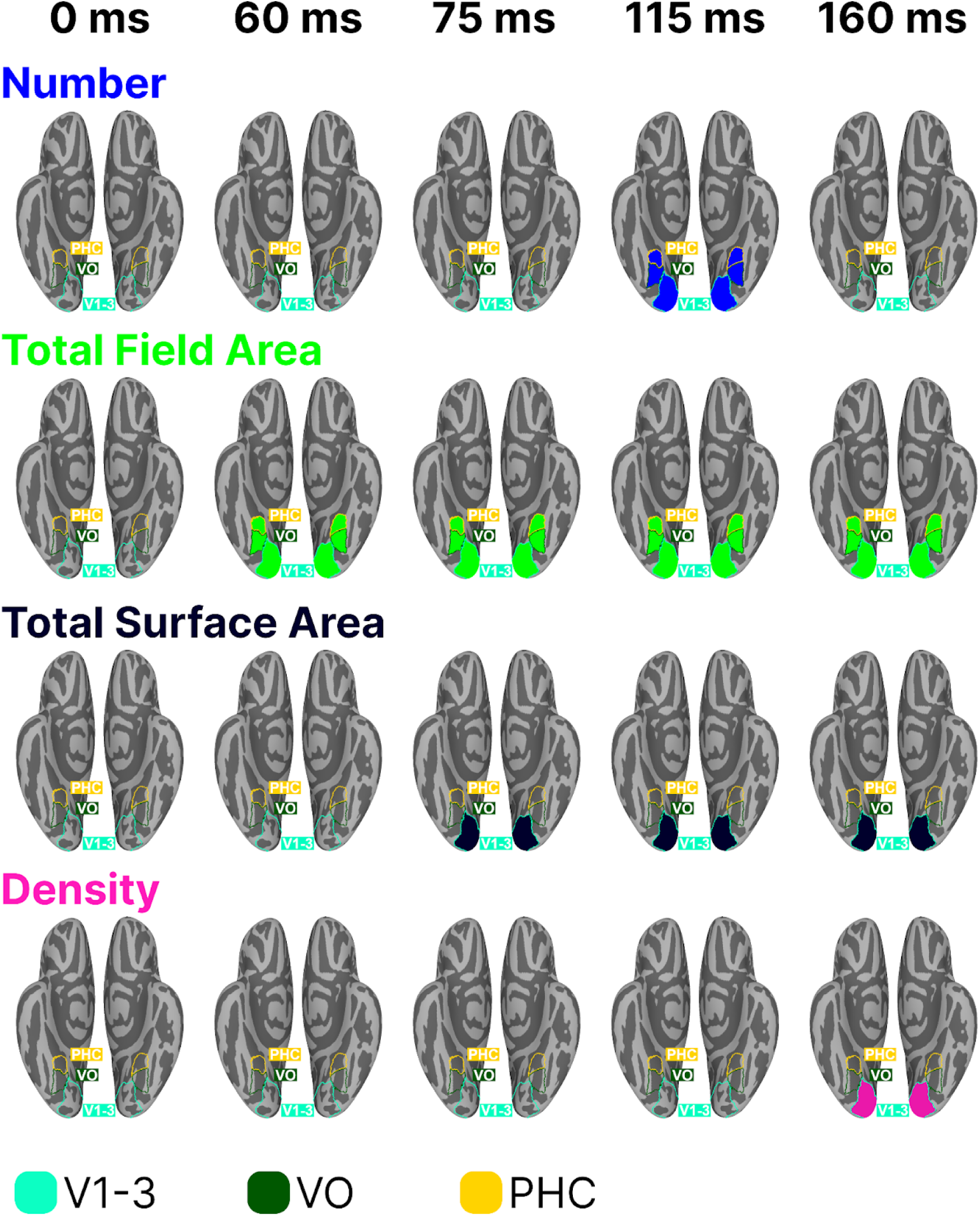
Spatiotemporal dynamics of four features (number, total field area, total surface area, and density) in the ventral stream regions are shown at five snapshots (time 0 ms when the stimulus begins, time 60 ms when total field area becomes significant, time 75 ms when total surface area becomes significant, time 115 ms when number becomes significant, and 160 ms when density becomes significant) from Videos 1 to 4. It’s important to note that the average item area is not depicted, as it did not show significance in any region.

## Discussion

This study investigated how and when the human brain encodes numerosity and non-numeric features in visual dot arrays. Using magnetoencephalography (MEG) combined with representational similarity analysis (RSA), semipartial correlations, MEG–fMRI fusion, and multidimensional scaling (MDS), we tracked the temporal evolution and cortical distribution of feature representations. This multivariate, time-resolved approach allowed us to disentangle the unique neural signatures of numerosity from correlated visual properties and to visualize how their representational geometry transformed over time. Together, these analyses tested 3 main questions:

1. whether number constitutes a primary perceptual attribute,
2. whether its encoding follows a hierarchical progression from early visual to associative areas,
3. Whether and how its representational format dynamically evolves over time.

### 1. Number is encoded early, directly, and on a separate channel: it’s a primary visual feature

Using representational similarity analysis, we unfolded the temporal dynamics of numeric and non-numeric features in the stimuli. Numerosity emerged very early, before the pairs of non-numeric features from which it could be indirectly computed (total field area and density on one side; total surface area and average item area on the other).

This finding suggests that the visual system can access numerosity information without first deriving it from these non-numeric features, and thus that it can be considered a primary visual property of the image, as first suggested by Burr and Ross (2008). A second line of evidence strengthening this claim is that numerosity was encoded in a distinct frequency band compared to all non-numeric features.

Neural oscillations are known to structure visual and cognitive processing in frequency-specific ways (Ward, 2003). In vision, gamma rhythms support feature binding (Fries, 2009), whereas theta oscillations contribute to processes such as spatial navigation (Buzsáki & Moser, 2013). In language, frequency bands dissociate cognitive operations: theta tracks lexical retrieval, alpha reflects semantic composition, beta supports sentence-level integration, and gamma indexes predictive processes (Lam et al., 2016). Across domains, different frequency channels appear to route distinct computational operations. Consistent with previous findings (Rubinsten et al., 2020), we observed non-symbolic numerosity representations primarily in the lower beta band, whereas density and total surface area shifted toward higher beta frequencies. This separation suggests the brain may minimize interference by allocating numerical and non-numerical processing to distinct oscillatory channels, a mechanism compatible with models proposing that different oscillatory regimes reflect distinct neurophysiological mechanisms (Cannon et al., 2013).

To the best of our knowledge, prior to our study, RSA was adopted to investigate the timing of numerosity encoding in the brain only by Bankson et al. (2019). However, their study encountered two significant limitations that complicate the interpretation of their results. Firstly, although they attempted to minimize the influence of non-numeric features on number prediction, they did not account for these effects when correlating neural RDM with number RDM, a step we rigorously followed in this research. Secondly, their selection of numbers did not ensure equal perceptual discriminability, as we did in our study. This raises the possibility that, given our uncertainty about whether these numbers are perceptually discriminable, the observed differences in activation across numbers could be influenced by other non-numeric features that can be perceptually discriminated more easily.

In contrast to our finding that numerosity uniquely explains the variance of MEG signals over other non-numeric features, Gebuis & Reynvoet (2013) reported no numerosity-related effects in the ERP signals of EEG during both passive and active viewing of a visual set of dots. This absence could stem from several factors, including the smaller sample size of their study, but also from their stimulus design, wherein they introduced a non-linear relationship between numerosity and non-numeric features in an attempt to de-correlate them. This design choice led to a higher variance in the non-numeric features than in number. This could potentially mask the effect of number in the ERP signals. Our findings do fit with two previous studies that used univariate ERP signals in EEG analysis to explore the timing of number representation (Park et al., 2015; Fornaciai et al., 2017). Paralleling our observations, their results indicate a rapid and direct encoding of numerosity information. However, a direct comparison between the timing of our results and these two studies is not straightforward, given that we designed our stimuli and analyzed the data differently. We employed perceptually definable non-numeric features to create our stimuli, while they used two orthogonal mathematically defined constructs (size and spacing) to define theirs (DeWind et al., 2015). Additionally, we ensured that number and average item area were chosen to be equally discriminable, and that the total field areas were sparse enough to emphasize numerosity processing, yet still allowed participants to reliably estimate number rather than rely on density cues (Anobile et al., 2013), whereas (Park et al., 2015; Fornaciai et al., 2017) did not control for the perceptual discriminability of size and spacing since these constructs were mathematical and inherently non-perceptual. In addition to our distinct approach in stimulus design, the current study also differed from the previous ones in data analysis. Whereas previous studies typically controlled only a subset of non-numeric visual features (e.g., size and spacing), leaving open the possibility that early numerical signals reflected other correlated dimensions, the present study included all four canonical non-numeric dimensions—total field area, density, total surface area, and average item area—as covariates in a semipartial correlation analysis. This allowed us to isolate variance uniquely attributable to numerosity from variance explained by each non-numeric feature. Consequently, the effect we reported for numbers reflects its unique influence, with all other non-numeric features having been partialled out.

Importantly, when we performed source reconstruction (see supplementary material), the onset of number representation appeared slightly earlier than in the sensor-level analysis, emerging around ∼75 ms. This timing is more consistent with the early effects reported by Park et al. (2015). Source reconstruction helps reduce the spatial blurring inherent in sensor measurements, allowing a clearer estimate of when specific cortical regions become engaged (Edelman et al., 2015; Wen et al., 2019). Notably, the spatial pattern of these source-localized representations aligns well with the RSA ROI results from our previous fMRI study (Karami et al., 2025), though the MEG patterns are somewhat more spatially diffuse, which is expected given the limits of MEG source estimation.

To further characterize how these representations unfold over time, we performed time generalization analysis, which assesses whether the representational format is stable (generalizing across time) or rapidly evolving. Numerosity showed a sustained and temporally generalizable pattern, consistent with a stable representational code. All non-numeric features displayed similar stable patterns, except average item area. This feature produced only a short-lived, highly dynamic pattern. These dynamics resemble intracranial findings showing distributed, rapidly evolving codes in high-level cortex (Rogers et al., 2021). Because fMRI temporally averages neural activity, such dynamic or sign-reversing codes are likely to be washed out, explaining why prior fMRI studies (Castaldi et al., 2019; Karami et al., 2025) did not detect average item area representations: the representation exists, but only briefly and in a rapidly changing format.

### 2. Number is encoded along the dorsal and ventral stream extremely fast and with no detectable temporal hierarchy

Previous studies, employing a regression-based approach, have revealed that a significant portion of the variance in the ERP can be explained by numerosity very early in time, approximately around 75 ms (Park et al., 2015) and 90 ms (Fornaciai et al., 2017) post-stimulus on the medial occipital channel. Based on these findings, it had been concluded that the rapid encoding of numerosity information initiates from the early visual cortex. However, their conclusions were primarily based on the examination of scalp topographies. Relating scalp topographies to underlying source locations is challenging due to the well-established phenomenon in which electrical potentials from various sources blend at the level of scalp EEG recordings (Baillet, 2017). Subsequent studies (Park, 2018; Guillaumé et al., 2018; Lucero et al., 2020; Van Rinsveld et al., 2020) sought to strengthen these initial findings by using steady-state visual evoked potentials (SSVEP). The rationale behind this method is to present stimuli flickering at specific frequencies, thereby tagging the neural activity pattern in the brain with the desired stimulus frequency (Vialatte et al., 2010; Norcia et al., 2015). Due to their robustness against artifacts and high signal-to-noise ratio, these evoked potentials are suitable for isolating responses to numerosity in both children and adults; however, they do not contain temporal information. All of the aforementioned studies, including those conducted by Park (2018) in children and Guillaumé et al. (2018), Lucero et al. (2020), and Van Rinsveld et al. (2020) in adults, provide evidence that SSVEPs of the occipital channels of EEG are modulated by numerosity. However, their claim that the early visual cortex is the source of activity is unjustified as it relies solely on the scalp topography of the EEG response, without any source reconstruction to localize the anatomical sources of signals (Baillet, 2017). Indeed, Van Rinsveld et al. (2021) recently enhanced the initial SSVEP paradigm (Van Rinsveld et al., 2020) with source localization, and the results suggest the presence of multiple cortical sources, including the IPS, supplementary motor area, and middle temporal gyrus.

Here, through the use of model-based MEG-fMRI fusion, we were able to unravel the temporal dynamics of numeric and non-numeric features within various brain regions along both the dorsal and ventral streams. Our findings indicate that numerical information is rapidly encoded along both the dorsal and ventral streams, preceding the pairs of non-numeric features (total field area-density, and total surface area-average item area), the combination of which could be used to compute number. One possible interpretation of these findings is that, akin to the operation of sensory maps (Young, 1998), where multiple repetitions of topographically organized maps operate parallelly to analyze various aspects of sensory input, different cortical regions in the brain may work in parallel to analyze numerical information. Interestingly, as demonstrated in our previous fMRI study (Karami et al., 2025) using searchlight analysis, the representation of numerosity information is widely distributed across the cortex along both the dorsal and ventral streams, a wide-spread network that is strikingly similar to the network of numerotopic maps discovered by Harvey & Dumoulin, 2017. The distribution of numerosity information across various brain regions, including those typically associated with different functions such as mental arithmetic, decision-making, motion processing, object processing, action planning, etc may suggest the use of numerosity information for diverse functions. This distributed architecture may also serve rapid recognition of numerosity information (Behrmann & Plaut, 2013). The idea of parallel pathways for number can be further supported by evidence of fast subcortical routes in vision. In primates, a motion-selective area MT/V5 receives a direct koniocellular projection from LGN or pulvinar, bypassing V1 and enabling extra-striate activation with minimal delay (Sincich et al., 2004; Bridge et al., 2015). By analogy, one can speculate that numerosity-selective regions beyond V1 might also receive unusually early subcortical inputs. In fact, a recent computational study of mouse LGN demonstrated that its responses carry information about numerosity alongside basic visual features (Gamal & Eldawlatly, 2025), implying that even the earliest stages of the visual hierarchy are sensitive to object count. In line with this idea, evidence from behavioural monocular–dichoptic paradigms, which suggest subcortical involvement (Collins et al., 2017) and from 7T fMRI demonstrating tuned responses to haptic numerosity in the putamen (Hofstetter & Dumoulin, 2021) supports the contribution of subcortical structures to numerosity processing, though their precise functions remain unclear.

Complementing these anatomical and modeling insights, intracranial recordings during arithmetic judgment tasks show that activation propagates through a chain of cortical “math” sites with astonishing speed. Pinheiro-Chagas et al. (2024) measured high-frequency broadband onsets from nine regions of interest, spanning ventral temporal (fusiform, inferior temporal gyri), dorsal parietal (IPS, SPL), and frontal cortices, as participants evaluated both simultaneous (“48 + 5 = 57”) and sequential (“4” “+” “8” “=” “5” “7”) addition problems. They found a highly linear progression of response-onset latencies (ROLs), with each successive region engaging on average 13.3 ms (SD = 11.3 ms) after the previous one, placing parietal activation roughly 250 ms after stimulus onset, i.e., about 100 ms after the initial fusiform response.

Following these observations, an alternative interpretation is an extremely fast but hierarchical propagation of numerosity from early visual cortex to associative regions. In this view the encoding of numerosity must be abstracted from local spatial representations of the visual input to form a global code, a transformation that appears to manifest along the intraparietal sulcus (Viswanathan & Nieder, 2020). Consistent with a hierarchical sequence, Nieder & Miller (2004) reported that numerosity-selective neurons in the fundus of the intraparietal sulcus (F-IPS) began responding at ∼73 ms after stimulus onset and conveyed numerosity information at around 106 ms, whereas prefrontal neurons (LPFC) reached numerosity selectivity at ∼165 ms. Thus, the parietal cortex led the prefrontal cortex by approximately 27–59 ms. However, such small latency offsets between early visual and higher areas are difficult to resolve noninvasively. Population-level time-generalization work shows that representations that remain stable over time can be easily read out by other brain areas, because the same neural pattern continues to carry the information without requiring precise timing or dynamic transformations; broad temporal generalization therefore signals a readily accessible code that downstream areas can exploit (Murray et al., 2016). Consequently, rapid hierarchical propagation (with small, hard-to-detect latency differences) and temporally stable population formats are complementary explanations: a stable/accessible code could enable fast readout by later stages even when interregional delays are only a few tens of milliseconds, and our MEG analyses may lack sensitivity to dissociate those tiny offsets.

While model-based MEG–fMRI fusion offers a powerful way to relate the temporal resolution of MEG to the spatial specificity of fMRI, it is important to note that the two modalities measure neural activity at different spatial scales (Khaligh-Razavi et al., 2018; Cichy & Oliva, 2020). Sensor-level MEG patterns reflect combined activity from distributed cortical generators, whereas fMRI ROI patterns index responses within more localized populations. As a result, fusion results should be interpreted as shared representational structure rather than precise one-to-one localization across modalities. In addition, representational similarity across regions can introduce interpretational ambiguity when their RDMs are correlated (See Figure S3 in the supplementary material).

To further examine the temporal progression and spatial distribution of numerosity representations, we therefore complemented the fusion results with time-resolved RSA and MDS applied to source-reconstructed MEG data. These analyses (Figure S4 and S5 in the supplementary material) supported the fusion findings, indicating rapid numerosity-related responses across retinotopically organized areas along both the dorsal and ventral streams. However, the absence of clear temporal separation between these regions should be interpreted with caution, as limitations of source reconstruction—including spatial leakage and smoothing—can obscure fine-grained temporal differences (Palva et al., 2018).

### 3. The neural representational geometry of number dynamically evolves

Results from multidimensional scaling (MDS) applied to the time-resolved representational dissimilarity matrices (RDMs) revealed a two-stage temporal profile of numerosity processing, suggesting that distinct coding schemes emerge over time. The first stage appeared around 115 ms after stimulus onset, followed by a second stage at approximately 220 ms. At this later stage, the MDS configurations showed the emergence of a curved representational structure that persisted across several subsequent time points (e.g., 230, 310, and 315 ms), closely resembling the geometry observed in our previous fMRI study (Karami et al., 2025). In that study, early visual regions exhibited a linear “number line” consistent with a monotonic summation code, whereas associative regions, particularly the intraparietal sulcus (IPS) and ventral occipitotemporal cortex, displayed a curved structure in which mid-range numerosities were separated from extremes. Such curved structures have been linked to numerosity-tuned neural populations, whose Gaussian-like response profiles peak at preferred numerosities (Piazza et al., 2004; Paul et al., 2020), while linear layouts are characteristic of monotonic codes, where neural activity systematically increases or decreases with numerosity. Simulations from Karami et al. (2025) further support this interpretation: monotonic response yields straight MDS configurations, whereas tuned response produces curved ones, reflecting a temporal transition from monotonic to tuned representational geometry. While these simulations do not conclusively establish the presence of tuned representations, they suggest that changes in response profiles over time could alter the global structure of the representational space.

The emergence of curved manifolds in the present MDS analyses thus likely reflects a temporal transformation in the format of numerosity coding, from an early monotonic response toward a later, tuned response. Importantly, this transformation should not be interpreted as a sequential progression across distinct cortical regions. Although our fMRI results localized linear and curved structures predominantly to early visual and associative regions respectively, monotonic responses have also been observed in associative cortices (Paul et al., 2020). Therefore, we interpret these temporal dynamics as reflecting changes in the geometry of population activity over time, rather than evidence for a strict hierarchical feedforward sequence across the dorsal stream.

This temporal evolution of numerosity representation parallels previous findings in both infants (Gennari et al., 2023) and adults (Park et al., 2015; Fornaciai et al., 2017), which similarly point to multiple stages of processing. One hypothesis proposes that numerosity perception unfolds in two stages: an initial encoding of raw numerosity, followed by a later stage supporting conscious numerical perception. Fornaciai and Park (2018) reported a connectedness effect emerging around 150 ms in EEG and corresponding discriminable patterns in area V3 with fMRI, suggesting that this later activity reflects segmentation into perceptual units. Subsequent work using masking and serial-dependence paradigms (Fornaciai & Park, 2021) showed that these connectedness-driven biases operate on perceived rather than physical numerosity, even when feedback and awareness are suppressed, supporting the interpretation that the ∼150 ms stage indexes perceptual numerosity. Although our current data do not directly measure conscious awareness, previous studies indicate that ERP correlates of visual awareness typically appear between 170–290 ms after stimulus onset (Dembski et al., 2021). Accordingly, the dominance of numerosity-tuned responses in this later period, likely involving parietal associative regions, may correspond to the emergence of conscious number perception and behavioral discrimination.

In summary, using MEG and MEG/fMRI fusion we revealed that numerosity representations emerge rapidly after stimulus onset, earlier than and independent from pairs of non-numeric features that could otherwise define it, indicating that numerosity functions as a primary perceptual feature. Furthermore, numerical representations arise swiftly across retinotopic regions of both the dorsal and ventral streams. These representations appeared prior to the encoding of non-numeric feature pairs and did not follow a strictly hierarchical temporal sequence, suggesting a simultaneous, distributed emergence of numerical information across visual pathways. Finally, multidimensional scaling (MDS) revealed a clear evolution in representational geometry over time, from an early, linear organization consistent with a summation-like code in occipital regions to a later, curved geometry indicative of numerosity-tuned coding in higher-order dorsal areas.

## Materials and Methods

### Participants

Thirty-nine healthy adults (twenty-one females, average age 25.1 years) with either normal vision or vision corrected participated. The University of Trento’s ethics committee in Italy approved the study, and all participants provided written informed consent and were compensated for their participation. To keep head movement below 10 mm, data from six participants were omitted from the final analysis. Additionally, three more participants were excluded because of poor behavioral performance (accuracy below 60%). This resulted in a final sample of thirty subjects (17 females, average age 25.6 years).

### Stimuli and procedure

Before the MEG experiment, participants were introduced to the task through 20 practice trials conducted outside the MEG environment. During the MEG scanning session, participants were presented with arrays of black dots on a mid-gray background. These dots orthogonally varied in terms of number, average item area, and total field area. There were 32 conditions resulting from the combination of four numerosities (six, ten, seventeen, or twenty-nine dots), four average item areas (0.04, 0.07, 0.12, 0.2 visual square degrees), and two total field areas (defined by a virtual circle with diameters of approximately 9 or 13.5 visual degrees; Figure 1). The selection of these numbers and average item areas was based on their perceptual discriminability, as established in previous research (Castaldi et al., 2018; Castaldi et al., 2019). The total field areas were chosen to ensure suitably sparse arrays of dots (1 dot/vd^2^) to be within the numerosity estimation regime and not within the density estimation regime (Anobile et al., 2013).

In each trial, a set of dots was presented for 500 ms and displayed over a wide thin red fixation cross (the sample set). Participants were asked to estimate the number of dots and hold that information in memory until the appearance of the next set, after a variable interstimulus interval (ISI) of 500 or 800 milliseconds (Figure 1). When the color of the fixation cross changed from red to green, participants were instructed to compare the number of dots in the current set (the match set) with the previous set and indicate whether it was larger or smaller by pressing one of two buttons, following provided instructions. After a delay of 1.8 seconds, the next trial began. Match sets differed by approximately 2 just noticeable differences (JNDs) in numerosity from the previous sample stimulus, based on the average numerosity JND estimated from a separate group of healthy adults (Castaldi et al., 2018). The other features (total field area and average item area) remained constant. Match trials occurred approximately 20% of the time.

The experiment consisted of twelve runs, each containing four blocks. Each block included thirty-six trials, comprising four match trials and thirty-two sample trials, one for each of the thirty-two conditions (4 number × 4 average item area × 2 total field area). After the sixth run participants’ response assignments were switched. Brief practice sessions were conducted at the beginning of the experiment and after the hand assignment change. Each run had a duration of approximately 3.5 minutes.

### MEG recordings and preprocessing

Subjects’ brain activity was recorded at the Center for Mind/Brain Sciences at the University of Trento using a MEG system comprising 306 channels (204 planar gradiometers and 102 magnetometers) manufactured by Elekta-Neuromag Ltd. in Helsinki, Finland. The data was acquired at a sampling rate of 1000 Hz and underwent online band-pass filtering within the frequency range of 0.01–330 Hz. The participants were seated upright within a room that was shielded against magnetic interference (AK3B, Vakuumschmelze, Hanau, Germany). Prior to the recording session, the unique head shape of each participant was measured using a Polhemus FASTRAK 3D digitizer (Polhemus, Vermont, USA). This process involved acquiring three fiducial points (nasion and pre-auricular points), five head position indicator (HPI) coils (one on each of the left and right mastoids and three on the forehead), and approximately 100 more locations distributed across the subjects’ skull. At the start of each run, head positioning inside the MEG helmet was measured by inducing a non-invasive current through the HPI coils.

Additionally, both horizontal and vertical electro-oculograms (EOGs) were recorded, along with an electrocardiogram (ECG), with the intention of later offline eliminating artifacts related to eye movements and heart activity. For stimulus presentation and control, Psychtoolbox 3 (Brainard, 1997) was employed. Each visual stimulus image was rear-projected onto a screen positioned at a distance of 120 cm from the participant’s eyes using a VPixx PROPixx projector.

The offline raw MEG data underwent a visual inspection to identify and eliminate noisy channels. Subsequently, we employed the MaxMove function of the Elekta Maxfilter software to remove head motion and to denoise the data using Maxfilter Signal Space Separation (Taulu et al., 2003). Following the maxfiltering, we proceeded with additional preprocessing using MNE-Python version 0.24.1 (Gramfort, 2013). First, the data underwent band-pass filtering within the range of 0.05 to 330 Hz and bandstop filtering at 50 Hz and its harmonics for line noise removal. To eliminate artifacts caused by eye movement and heart rate, independent components of the MEG data were computed and correlated with the EOG and ECG signals. Components displaying significant correlations were subsequently removed through manual inspection. Subsequently, to enhance signal-to-noise ratio (SNR) and lower computational demands, we applied temporal smoothing on signals using Savitzky–Golay filter (with order 4 and frame size 25; Acharya et al., 2016) and downsample the data to 200 Hz (Grootswagers et al., 2017). Following these preprocessing steps, the data was epoched into 0.9-second trials, starting 100 ms before the stimulus onset. Each epoch was then normalized by subtracting the baseline period mean. Given that magnetometer and gradiometer sensors have different measurement units, we opted to exclusively use gradiometer data for our subsequent analysis. This choice aligns with prior studies, such as Ritchie et al. (2015), which exclusively focused on gradiometer measurements, as well as studies like Kaiser et al. (2016) that concentrated solely on magnetometer data. Additionally, some studies, such as Proklova et al. (2019), have chosen to report results separately for each sensor type.

### Time Resolved Multivariate Representational Similarity Analysis (RSA)

To explore whether and when the representations of numeric and non-numeric features of the stimuli can be disentangled from the brain signal we employed representational similarity analysis combined with semipartial correlation (Kriegeskorte et al., 2008; Kriegeskorte & Kievit, 2013).

To create neural representational dissimilarity matrices (RDMs), we organized the preprocessed MEG data into pattern vectors, each encompassing MEG data from all selected channels for each trial and time point. Following this, we averaged data from trials of the same conditions into single trial representations. Consequently, this process yielded 32 distinct MEG patterns, each corresponding to one of the 32 conditions, for each time point (100 ms before to 600 ms after stimulus onset). To quantify the similarity between all pairs of these 32 patterns, we employed Pearson correlation. We then subtracted these correlation values from 1 to generate 32 x 32 MEG RDMs for each time point. Following the approach outlined in our previous work (Karami et al., 2025), we then conducted a semipartial correlation analysis (using Pearson correlation) to assess the extent to which the dissimilarity structure in MEG patterns could be attributed to multiple predictor or model matrices that capture key quantitative dimensions of the stimuli. These dimensions include number, average item area, total field area, total surface area, and density. These five model RDMs quantitatively represent the logarithmic distance between pairs of stimuli in terms of number, average item area, total field area, total surface area, and density. The semipartial correlation between a vectorized neural RDM and the specific model RDM under examination measures the unique variance shared between the neural RDM and the chosen model RDM while partialling out the effect of all other models RDM from the neural RDM. A diagram outlining this procedure is presented in Figure 2. The analysis was implemented using the CoSMoMVPA MATLAB toolbox (Oosterhof et al., 2016) and custom-written code in MATLAB R2019 (The MathWorks, Inc., Natick, MA).

### Sensor-space Searchlight Multivariate RSA

In order to identify sensors over time where the independent contribution of five model RDMs to the neural RDM could be observed, we conducted a sensor-space searchlight analysis (Kriegeskorte et al., 2006; Proklova et al., 2019). Applying the method described in the preceding section, we employed semipartial correlation analysis (Pearson correlation) to evaluate the degree to which the dissimilarity structure in MEG patterns be explained by the model matrices. To achieve this, we designated a neighborhood for each MEG channel comprising its 20 nearest MEG channels. Subsequently, we conducted time-resolved RSA for each MEG channel, with the analysis being restricted to the data from its neighboring channels. We proceeded to average the results in 150 ms bins, producing a single map representing the grand average correlation coefficient for each feature.

The analysis was carried out using the CoSMoMVPA MATLAB toolbox (Oosterhof et al., 2016) alongside custom-written code in MATLAB R2019 (The MathWorks, Inc., Natick, MA). The results were then visualized using the ft_topoplotER function in the FieldTrip Toolbox (Oostenveld et al., 2011).

### Time-Frequency Resolved Multivariate RSA

We employed time-frequency decomposition to investigate whether the representation of numbers occurs in a distinct frequency band compared to other non-numeric features. These findings could be crucial to indicate whether a number is encoded separately from other non-numeric features. Following Xie et al., (2022), we produced time-frequency decompositions of the preprocessed MEG time series using Morlet wavelets for each trial and sensor. The wavelets used in the analysis had a consistent length of 2,600 ms and were logarithmically distributed across 40 frequency bins ranging from 2 Hz to 30 Hz. Absolute power values for every time point and frequency bin were derived by taking the square root of the resulting time-frequency coefficients. These power values were then normalized to represent relative changes, expressed in decibels (dB), concerning the pre-stimulus baseline (–100 ms to 0 ms in relation to stimulus onset). This analysis was carried out using custom-written MATLAB R2019 code (The MathWorks, Inc., Natick, MA). We decompose the MEG time series using a custom function developed by Xie et al., (2022; available at https://github.com/siyingxie/VCR_infant/blob/main/code/timefrexdecomp.m).

### Temporal Generalization Analysis

We employed a temporal generalization analysis (King & Dehaene, 2014) to examine how the neural representation of numbers and related features evolves over time. This method goes beyond testing whether a feature is decodable at a single latency: it assesses whether a classifier trained at one time point can successfully decode the same feature at other time points. Sustained cross-temporal decoding indicates that the underlying representational format remains relatively stable, whereas a strong diagonal with little off-diagonal generalization suggests that the neural code is rapidly transforming across time, a diagnostic signature of dynamic coding also highlighted by Rogers et al. (2021).

To implement this analysis, We first randomly combined every four trials into pseudo-trials in order to enhance the overall signal-to-noise ratio (Guggenmos et al., 2018). Subsequently, we trained a support vector machine (SVM) classifier to decode our four numbers (6, 10, 17, and 29), two total field areas (9, 13.5), four average item area (0.04, 0.07, 0.12, 0.2), sixteen total surface areas (resulting from multiplying four different numbers by four different average item areas), and eight different densities (resulting from dividing four different numbers by two different total field areas) at a specific time point and then tested the same classifier across all other time points separately. The preceding steps were repeated for a total of twenty randomized assignments of trials to pseudo-trials (permutations). Finally, we computed the average decoding accuracy across all permutations. The analysis was carried out using the CoSMoMVPA MATLAB toolbox (Oosterhof et al., 2016) and LIBSVM (Chang and Lin, 2011).

### MEG-fMRI Fusion with RSA

To relate the temporal dynamics captured by MEG with the spatial patterns revealed by fMRI, we performed RSA-based MEG-fMRI fusion (Cichy et al., 2014; Cichy et al., 2016). This approach identifies when the representational structure in a given brain region, as measured by fMRI, corresponds to the representational structure derived from time-resolved MEG signals, indicating a shared representational format at that moment.

We computed Pearson correlation coefficients between each participant’s MEG representational dissimilarity matrix (RDM) and the group-averaged fMRI RDMs (N = 31) from predefined regions of interest, including early visual areas (V1–V3), dorsal regions (V3ABV7, IPS1–5), and ventral regions (VO, PHC). This analysis assessed the correspondence between the representational structures captured by MEG and those derived from fMRI across these regions. In our previous fMRI study (Karami et al., 2025), we used the same task as in the present experiment, differing only in stimulus timing, which was faster in the MEG version. Neural RDMs were computed from t-statistics of the first-level analysis by calculating correlation distances between activation patterns for all condition pairs within each ROI (V1–V3, V3ABV7, IPS1–5, VO, and PHC). Twenty-three participants overlapped between the two studies. Accordingly, we used the group-averaged fMRI RDMs as a robust estimate of the underlying pattern dissimilarity (Nili et al., 2014) for the MEG-fMRI RSA.

### Model-based MEG-fMRI Fusion

To unfold the temporal dynamics of different brain regions involved in processing numerosity and the non-numeric features we used model-based MEG-fMRI fusion (Cichy & Oliva, 2020). Here, we extended the MEG-fMRI fusion approach described in the previous section to incorporate multiple models into the fusion process using conjunction inference (Nichols et al., 2005; Khaligh-Razavi et al., 2018). This extension allowed us to spatiotemporally resolve the processing of both numeric and non-numeric information. The rationale behind this method is to combine the result from three different representational similarity analysis using a logical ‘AND’ operator: (I) MEG-to-model RSA, as detailed earlier; (II) fMRI-to-model RSA, as explained in our previous fMRI study (Karami et al., 2025); and (III) MEG-to-fMRI RSA, which resulted from Pearson correlation between MEG RDMs from individual subjects, as detailed earlier, and five group-averaged RDMs derived from thirty-one participants in five regions of interest (ROIs) as outlined in the Karami et al., 2025.

The result, employing the ’AND’ operator, highlights MEG time points associated with fMRI locations that have a similar representational geometry both to each other and to the model RDM of interest.

### Multidimensional Scaling (MDS)

While the results from RSA reveal how each feature contributes to the variance of our data in a hypothesis-driven manner, we also explore the latent structure of our data using a data-driven approach with MDS (Kruskal, 1964). This approach was applied directly to the RDM, and subsequently we visualized the first two dimensions of the MDS output. With MDS we visualize the organization of stimuli on a two-dimensional plot, where the distances between them accurately reflect the differences in the neural response patterns they evoked. Stimuli positioned closely together in these representations represent similar neural response patterns (Nili et al., 2014). The MDS analysis was conducted using the MATLAB function cmdscale on the group-average RDM across participants spanning various time points from the onset of stimulus presentation to 600 ms. The group-average RDM is calculated by averaging the RDMs from all thirty subjects. This group-level RDM is more accurate and less susceptible to noise compared to the RDMs obtained from individual subjects (Nili et al., 2014).

### Statistical Testing

Throughout this article, statistical significance was evaluated through a one-sample t-test against zero across subjects. To control for multiple comparisons, the results were corrected using threshold-free cluster enhancement (TFCE; Smith & Nichols, 2009), employing Monte Carlo simulations with 10,000 permutations, as implemented in the CoSMoMVPA MATLAB toolbox (Oosterhof et al., 2016). The resulting statistics were then thresholded at p < 0.01 (one-tailed) to determine significance. For the searchlight analysis, significance was evaluated independently for each time point using the TFCE method. This method helped identify sensors where the contribution of a specific predictor to neural dissimilarity was significantly above zero.

## Code accessibility

All the code related to this study is available through GitHub: https://github.com/alireza-kr/numerosity_fmri-meg-cnn/

## Declaration of generative AI and AI-assisted technologies in the writing process

The authors used ChatGPT to assist with rephrasing certain sentences during the preparation of this work. They subsequently reviewed and edited the content as needed and took full responsibility for the final published article.

## Acknowledgements

Thanks to Marco Buiatti for valuable comments and discussions. M.P. and A.K. acknowledge the Italian Ministry for Education, University and Research (MIUR) for the Departments of Excellence Grant supporting a PhD scholarship to AK and the Centre for Mind and Brain Sciences (CIMeC) for supporting the research. For this work, A.K. was awarded NENS Exchange Grants from Federation of European Neuroscience Societies (FENS) and the 42° European Workshop on Cognitive Neuropsychology (https://www.ewcn.eu/) prize. M.P. and E.C. were funded by the Italian Ministry of Education, University, and Research under the PRIN2022 program: M.P through the Project ‘Sense of number vs. sense of quantity: modeling, neuroimaging, behavior’, grant no.2022EBC78W; E.C. through the Project ‘RIGHTSTRESS—Tuning arousal for optimal perception’, grant no.2022CCPJ3J. A.K. and M.N.H. were supported by an Independent Research Group Grant to M.N.H. by the Max Planck Society. M.N.H. was supported by the ERC Starting Grant project COREDIM (ERC-StG-2021-101039712), a LOEWE Start professorship by the Hessian Ministry of Higher Education, Science, Research, and the Arts, and German Research Foundation (DFG) under Germany’s Excellence Strategy (EXC 3066/1 “The Adaptive Mind”, Project No. 533717223).

