## Supplementary File for "Numerosity Is Directly Sensed and Dynamically Transformed in the Human Brain: Evidence from MEG-MRI Fusion"

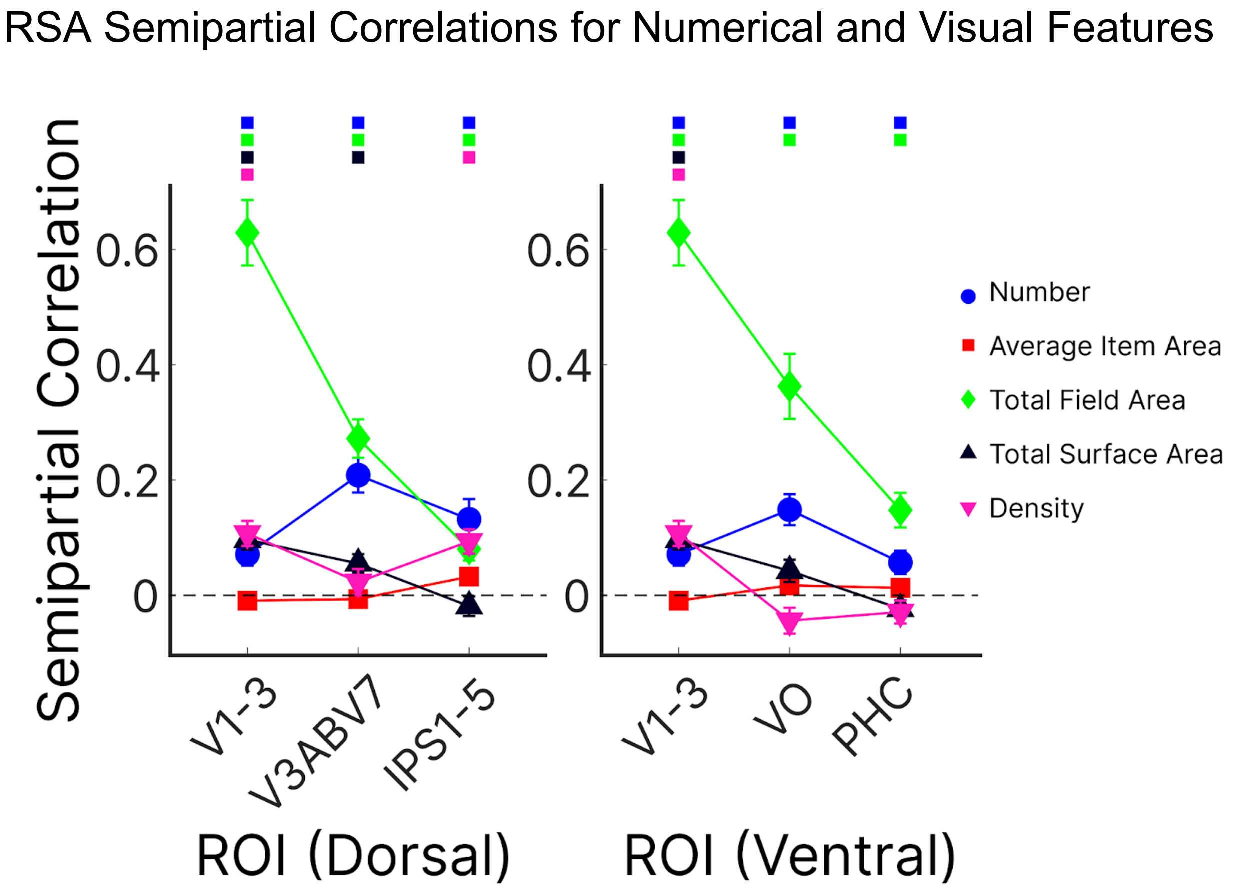


Figure S1. Semipartial correlation coefficient obtained from the representational similarity analysis for number, average item area, total field area, total surface area and density from predefined dorsal and ventral retinotopic ROIs. Data points show mean semipartial correlation coefficient across subjects (n = 31) ± standard error of the mean (SEM). The coloured points above the figure indicate where the effect significantly exceeds zero ($p<0.01$). Figure adapted from our previous fMRI study (Karami et al., 2025).

### MDS of Numerosity Representations Across Time and Modality

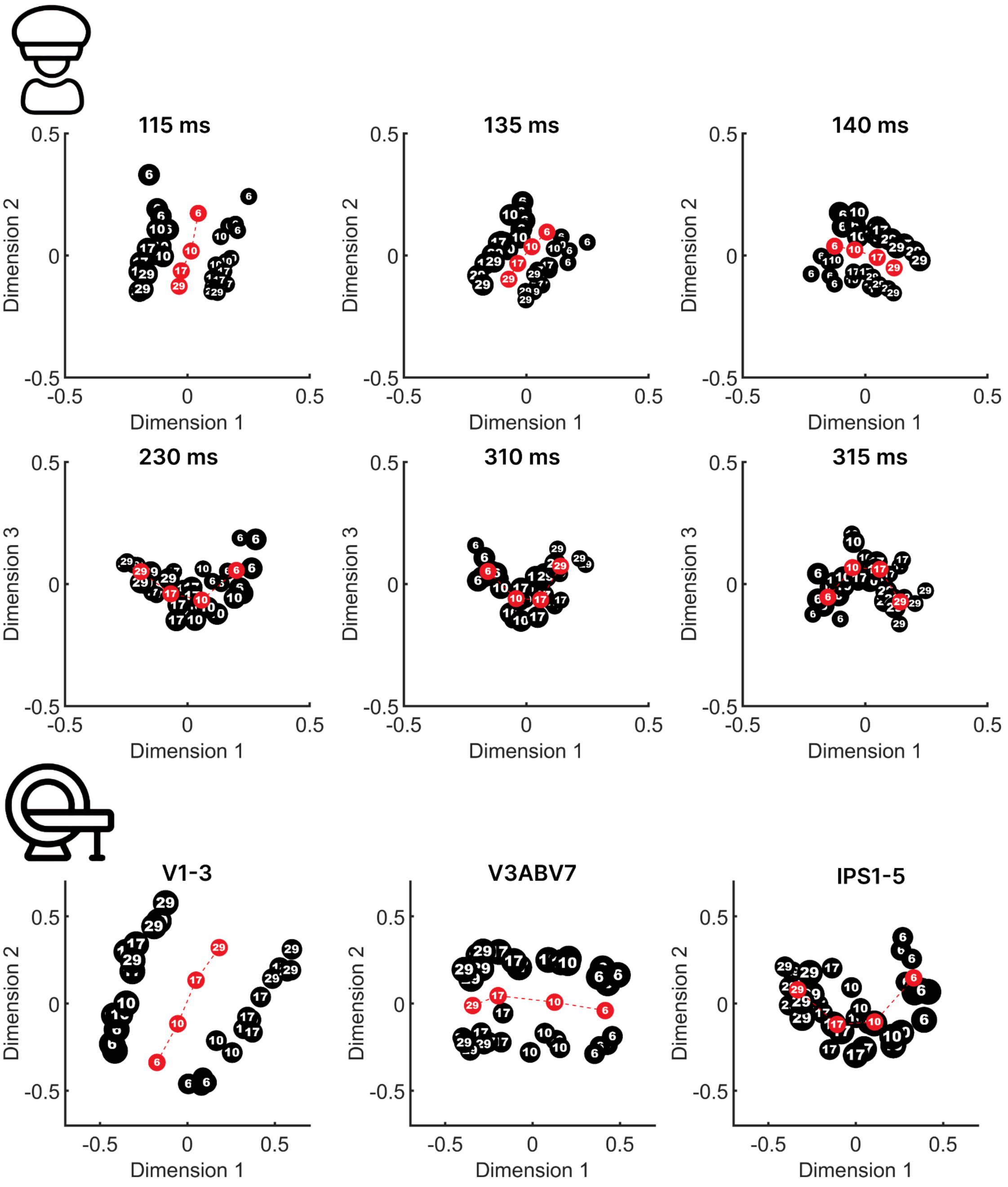


Figure S2. Multidimensional scaling (MDS) visualizes representational similarities between stimuli in a two-dimensional space at three early time points when numerosity information first becomes significant (115 ms, 135 ms, and 140 ms) and three later time points (230 ms, 310 ms, and 315 ms). The third row shows MDS results from our previous fMRI study for three key regions along the dorsal stream: V1–V3, V3ABV7, and IPS1–5. The black circles represent the 32 stimuli. The circle sizes vary, indicating stimuli with small total field area (small circles) and larger total field area (large circles). The red circles indicate the average coordinates of each number.

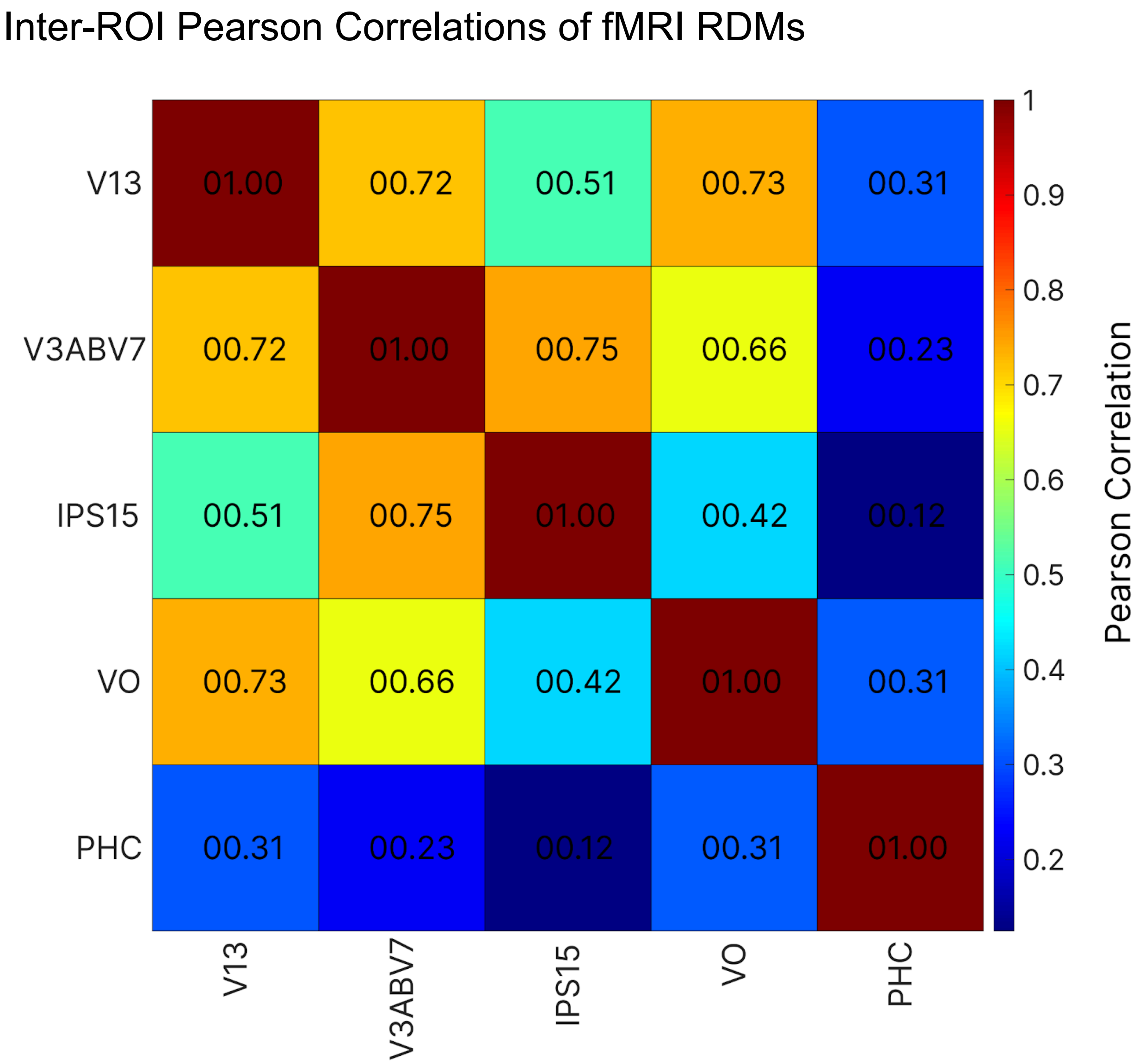


Figure S3. The Pearson correlation coefficient was computed across five group-average fMRI RDMs, involving 31 subjects. The regions of interest include early visual areas (V1-3), dorsal regions (V3ABV7, IPS1-5), and ventral regions (VO, PHC) as outlined in our previous fMRI study (Karami et al., 2025).

#### MEG Source Reconstruction

Following initial preprocessing outlined in the main text, we applied Minimum-Norm Estimates (MNE; Hämäläinen & Ilmoniemi, 1994) for cortical source estimation of MEG signals using the MNE-Python toolbox (Gramfort, 2013). Estimation of volume conduction was derived from the individual anatomies of the subjects, utilizing single-layer boundary element models (BEMs). These BEMs were generated for each subject's anatomy using the FreeSurfer software (Dale et al., 1999) and its watershed algorithm. The defined source space included 516 source points per hemisphere, located along the gray/white matter boundary as determined by FreeSurfer. Alignment of MEG/MRI data was achieved using fiducials and digitizer points on the head surface. The sensor noise covariance matrix was computed from the baseline period (−0.1 to 0 s relative to stimulus onset) and regularized following the Ledoit–Wolf procedure (Ledoit & Wolf, 2004). Source activations results were then projected on the SUMA standard mesh in AFNI (https://afni.nimh.nih.gov/) (Cox, 1996) using custom-written code in MATLAB R2019.

As Wen et al. (2019) show, decoding in the sensor space reflects a mixture of overlapping neural sources, where volume conduction and spatial blurring smear electrical activity across scalp sensors, effectively delaying or broadening the temporal profile of decoding peaks (Wen et al., 2019). When they localized activity to cortical ROIs (e.g., early visual cortex, LPFC), the reconstructed time courses revealed earlier and more distinct peaks of category- or relevance-based encoding—reflecting the underlying cortical dynamics more directly. Similarly, Edelman et al. (2016) demonstrated that EEG Source Imaging (ESI) mitigates the volume conduction effect by projecting scalp potentials back to their cortical generators, which reduces cross-talk from distant areas and increases spatial resolution (Edelman et al., 2015). This results in cleaner, more focal time courses that allow task-related neural responses (like motor or visual feature encoding) to emerge earlier and with higher discriminability than in sensor-level data. Consistent with these findings, in our study, the time-resolved RSA in source space revealed an earlier representation of numerical information compared to the sensor-space analysis, likely because source reconstruction disentangled overlapping generators and enhanced the detectability of early number-selective responses.

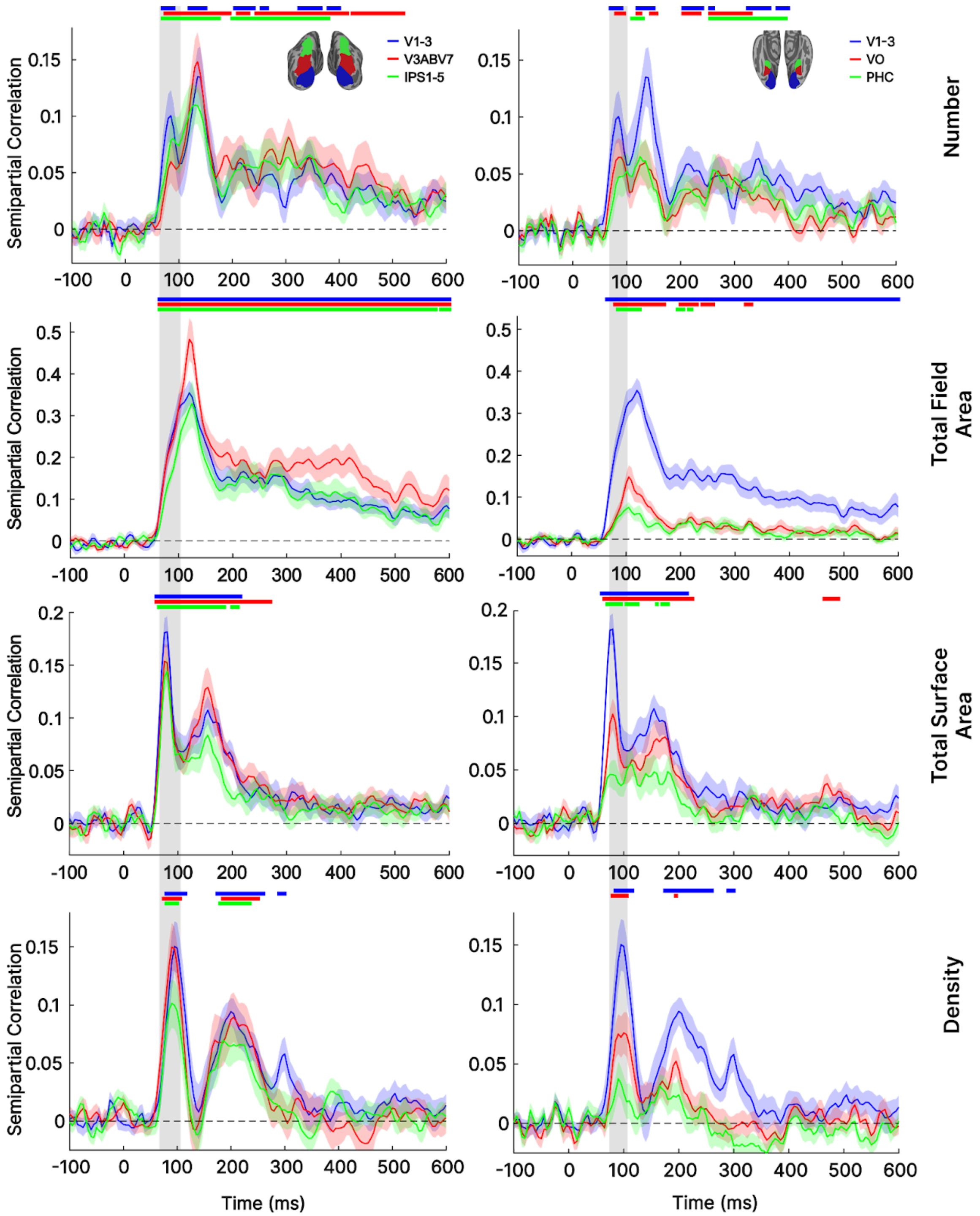


Figure S4. Semipartial correlation coefficients derived from the representational similarity analysis for number, total field area, total surface area, and density across various regions, including early visual areas (V1-3), dorsal regions (V3ABV7, IPS1-5), and ventral regions (VO, PHC), using source-reconstructed MEG. Standard error of the mean (SEM) across participants is depicted as a shaded area. The horizontal coloured dots indicate significant time points with effects significantly exceeding zero (thresholded at $p < 0.01$, TFCE corrected).

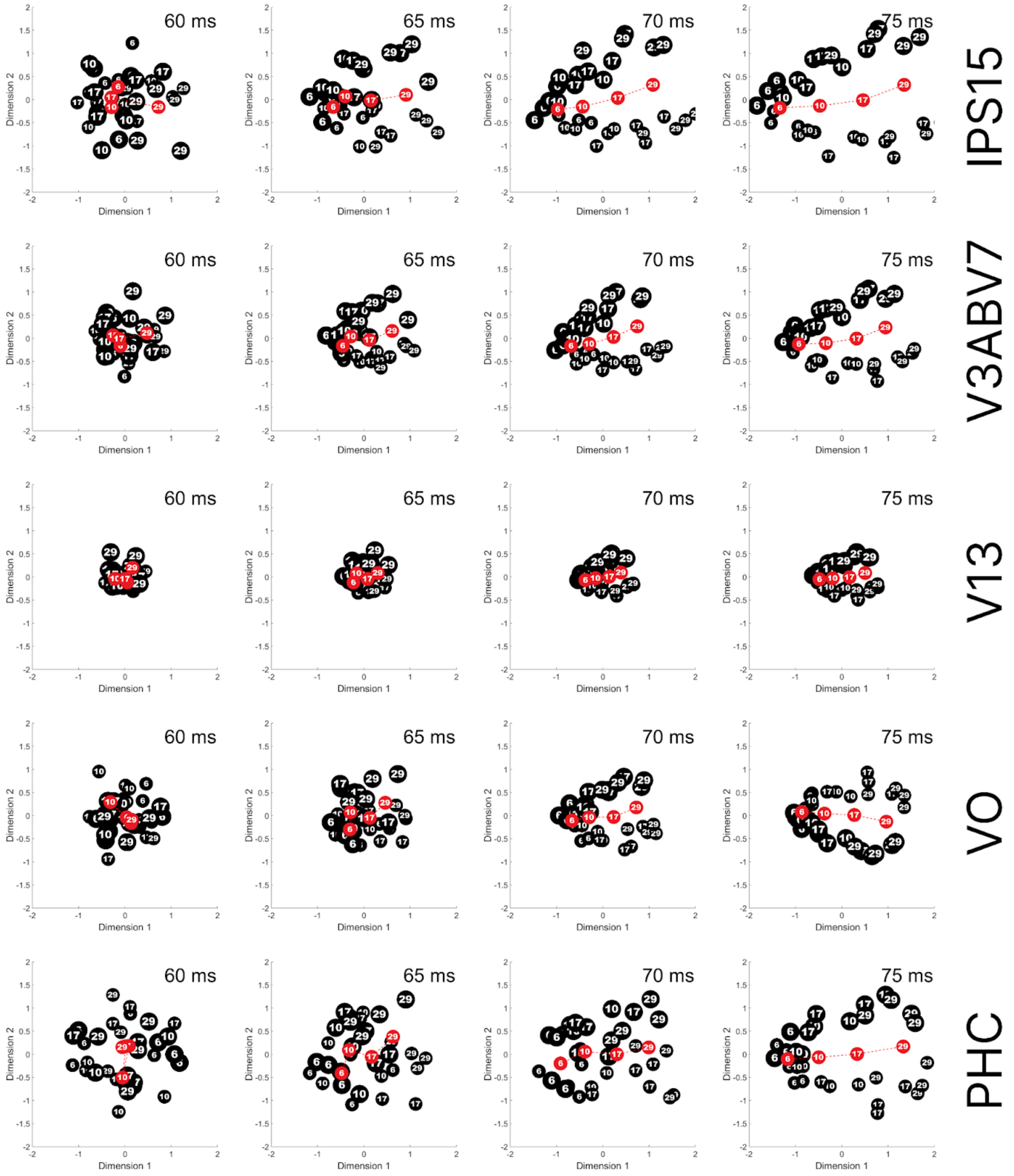


Figure S5. Multidimensional scaling (MDS) reveals representational similarities between stimuli in a two-dimensional space for four distinct time points: 60 ms, 65 ms, 70 ms, and 75 ms across various regions, including early visual areas (V1-3), dorsal regions (V3ABV7, IPS1-5), and ventral regions (VO, PHC), using source-reconstructed MEG. The black circles represent the 32 stimuli. The circle sizes vary, indicating stimuli with small total field area (small circles) and larger total field area (large circles). The red circles indicate the average coordinates of each number. The results reveal a clear rank-ordering of numbers, starting around 70 ms post-stimuli in all regions.
